# Phylogenomic evidence for host specialization and genetic divergence in OsHV-1 infecting *Magallana gigas* and *Ostrea edulis*

**DOI:** 10.1101/2023.08.23.554398

**Authors:** Camille Pelletier, Germain Chevignon, Nicole Faury, Isabelle Arzul, Céline Garcia, Bruno Chollet, Tristan Renault, Benjamin Morga, Maude Jacquot

## Abstract

Cross-species transmission is a major driver of disease emergence in humans and animals. The *Ostreavirus ostreidmalaco1* (OsHV-1) is mainly associated with mortality in the Pacific oyster *Magallana gigas*, but has also been found in other mollusks, including the European flat oyster *Ostrea edulis*. This raises questions about OsHV-1 host specificity. This study explored the genetic differentiation of OsHV-1 in *M. gigas* and *O. edulis* and the underlying mechanisms. Using high-throughput sequencing, 40 OsHV-1 genomes were obtained from both *O. edulis* and *M. gigas* and were analyzed to assess viral diversity, lineage isolation, and cross-species transmission. Comparative genomics, population genetics, phylogenetic and phylodynamic methods revealed that host species significantly influence viral genetic structure. The data suggest that OsHV-1 was introduced in Europe with *M. gigas*, followed by a cross-species transmission event and divergence into two distinct lineages. Selection signals were identified in genomic regions involved in key viral functions, including host binding, DNA replication, and membrane-associated proteins, indicating possible adaptation to different hosts. Future research should investigate coevolution between OsHV-1 and a broader range of host species using phylogenetic approaches to better understand host-virus dynamics.

## 1. Introduction

The study of multi-host-pathogen systems is crucial to improve our understanding of the spread and emergence of infectious diseases, and to develop effective strategies to address animal health problems. Many pathogens are able of multiplying themselves in various host species, and multiple pathogen lineages – defined as groups of genetically similar viruses with a common ancestor - may exhibit different phenotypic traits (Roche et al., 2013; Woolhouse et al., 2001). Pathogen lineage frequencies within host populations can vary depending on, for example, their transmission efficiency within and between host populations (Handel & Rohani, 2015; VanderWaal & Ezenwa, 2016), while barriers to transmission may result in population-specific patterns of infection for different lineages (Geoghegan et al., 2016; Kuiken et al., 2006; Park et al., 2013). Understanding the partitioning of pathogens genetic diversity among host species is particularly important for disease transmission when pathogens have the ability to infect both wildlife and livestock (Jones et al., 2008; Woolhouse et al., 2001). Disentangling epidemiological cycles in such systems is often challenging, yet investigating how the genetic diversity of pathogens is partitioned among host species is key to properly assess and manage disease risk (Archie et al., 2009; Roche et al., 2013).

Herpesviruses are an order of double-stranded DNA viruses commonly found across multiple species of vertebrates (*e.g.* fish, birds, pigs, horses, humans) and occasionally in invertebrates (*e.g.* mollusks). However, they are characterized by their narrow host range and are known to coevolve with their hosts, leading to a long-standing relationship between virus and host species (Azab et al., 2018; Bandín & Dopazo, 2011; Davison et al., 2005). Nevertheless, there have been several reports of cross-species transmission (CST) events in both vertebrates (*e.g.* transmission of cercopithecine herpesvirus 1 from primates to humans (Azab et al., 2018; Bandín & Dopazo, 2011; Huff & Barry, 2003)) and invertebrates such as mollusks (*e.g.* transmission of *Ostreavirus ostreidmalaco1* from Pacific oysters to scallops or flat oysters (Arzul, Renault, Lipart, et al., 2001; Azab et al., 2018; Bandín & Dopazo, 2011)). Herpesviruses affecting mollusks are restricted to the *Malacoherpesviridae* family which currently includes two species, the *Aurivirus haliotidmalaco1* (HaHV-1), formerly known as *Haliotid herpesvirus 1* infecting gastropods and the *Ostreavirus ostreidmalaco1* (OsHV-1), formerly known as *Ostreid Herpesvirus type 1* infecting bivalves. Several OsHV-1 lineages have been reported infecting a wide range of host species (Arzul, Nicolas, et al., 2001; Arzul, Renault, Lipart, et al., 2001; Arzul et al., 2017; Carnegie et al., 2016). Indeed, these lineages can infect and cause mortalities in many bivalve species, including the scallops *Pecten maximus* (Arzul, Nicolas, et al., 2001) and *Chlamys farreri* (C.-M. Bai et al., 2019), the clams *Anadara broughtonii* (Xia et al., 2015), *Ruditapes philippinarum* (Renault & Arzul, 2001; Renault, 1998, p. 199) and *Ruditapes decussatus* (Renault & Arzul, 2001; Renault, 1998), as well as the oysters *Magallana angulata* (Arzul, Renault, Lipart, et al., 2001), *Crassostrea virginica* (Farley et al., 1972), *Ostrea edulis* (Arzul, Renault, Lipart, et al., 2001; Comps, 1988; Renault et al., 2001) and *Magallana gigas* (Burioli et al., 2017; Morga-Jacquot et al., 2021; Nicolas et al., 1992; Renault et al., 1994).

In France, two oyster species have been shown to be infected by OsHV-1: the native European flat oyster *O. edulis* and the introduced Pacific oyster *M. gigas*. Pacific oysters were introduced in France at the end of the 1960s in response to the dramatic decline of the Portuguese oyster (*M. angulata*) populations (Grizel & Heral, 1991). *M. gi*gas is currently the most widely cultivated marine mollusk species in Europe and worldwide (FAO, 2023). While herpes like viruses were first observed in *C. virginica* adults in the United States of America in 1970, OsHV-1has been detected in *M. gigas*, in a hatchery in France, approximately 20 years after the introduction of *M. gigas* (Renault et al., 1994). OsHV-1 induces a major threat to juvenile Pacific oysters and has significant economic impacts in many production areas all around the world (C.-M. Bai et al., 2019; Burge et al., 2006; EFSA Panel on Animal Health and welfare (AHAW), 2010; Paul-Pont et al., 2013; Renault et al., 1994, 2012). On the other hand, currently OsHV-1 does not impact the flat oyster production, as it only leads to rare symptoms, in few cases accompanied by mortalities. Historically, *O. edulis* production was already compromised due to various pathogens infection, including *Marteilia refringens* and *Bonamia ostreae*.

Traditionally, OsHV-1 lineages have been characterized by targeted amplicon sequencing (Arzul, Nicolas, et al., 2001; Davison, 2002; Renault et al., 2001, 2012; Segarra et al., 2010). This approach allowed the characterization of OsHV-1 Var, a specific lineage detected in clams *R. philippinarum* in 1997 (Arzul, Renault, Lipart, et al., 2001) and scallops *P. maximus* in 2000 (Arzul, Nicolas, et al., 2001). This lineage has never been detected in wild clams or scallops, likely because only limited OsHV-1 surveillance has been conducted in these species in their natural environments. Targeted approaches also allowed the characterization of OsHV-1 µVar, the lineage mostly detected in Europe currently. This lineage was first detected in spring 2008 and has since been involved in mass mortalities of the Pacific oyster juveniles (Segarra et al., 2010).

According to these observations, and because herpesviruses generally have a limited ability to infect a wide range of hosts, the assumption of OsHV-1 host specialization cannot be ruled out (Arzul, Renault, Lipart, et al., 2001) despite the fact that several viral lineages have been detected sporadically in various bivalve host species worldwide (Arzul, Nicolas, et al., 2001; Arzul, Renault, & Lipart, 2001; Arzul, Renault, Lipart, et al., 2001; C. Bai et al., 2016; Renault et al., 2001). Indeed, the genetic information obtained so far suggests that different host species may be infected by distinct lineages of OsHV-1 (Morga-Jacquot et al., 2021; Xia et al., 2015), indicating an epidemiological cycle that involves a wide range of host species that still needs to be clarified.

One significant drawback of targeted approaches is that they cannot comprehensively capture the entire genome, leading to an incomplete interpretation of the genomic information (Maiden et al., 2013; Sunde et al., 2020). Advances in sequencing technologies allowed for the acquisition of the first OsHV-1 genome obtained from *M. gigas* larvae collected in 1995. This genome has since then been used as a reference genome (Davison et al., 2005). To date, more than hundred OsHV-1 additional genomes have been sequenced from four host species: *C. farreri*, *A. broughtonii*, *O. edulis* and *M. gigas* (Abbadi et al., 2018; C.-M. Bai et al., 2019; Burioli et al., 2017; Morga-Jacquot et al., 2021; Ren et al., 2013; Xia et al., 2015). The genome of the virus is linear and its size ranges from 199 kb to 211 kb. It contains from 123 to 128 predicted ORFs, of which only 57 have putative functions (Abbadi et al., 2018; C.-M. Bai et al., 2019; Burioli et al., 2017; Davison et al., 2005; Ren et al., 2013; Xia et al., 2015). OsHV-1 genomic architecture is similar to some other herpesvirus genomes (*i.e.* Herpesvirus group E such as HSV-1) with two invertible unique regions (U_L_ for Unique Long and U_S_ for Unique Short) surrounded by inverted repeats (TR_L_/IR_L_ and TR_S_/IR_S_). An additional unique sequence (called X) can be found between IR_L_ and IR_S_, resulting in the overall genomic organization that can be resumed as TR_L_-U_L_-IR_L_-X-IR_S_-U_S_-TR_S_ (Davison et al., 2005). Preliminary analyses of a few number of OsHV-1 genomes, isolated from different bivalve species, revealed a relatively high genotypic diversity (C.-M. Bai et al., 2019; Morga-Jacquot et al., 2021; Ren et al., 2013; Xia et al., 2015). More specifically, comparative genomic analyses of lineages infecting *O. edulis* and *M. gigas* showed the presence of two large deletion areas in the U_L_ region as well as host-specific polymorphisms along the genome (Morga-Jacquot et al., 2021). These results suggested that OsHV-1 evolution is strongly influenced by the host species (Morga-Jacquot et al., 2021). However, the low sampling size of previous studies did not allow the application of population genetic and phylodynamic analyses and still little is known about the level of genetic isolation of OsHV-1 lineages infecting *O. edulis* and *M. gigas* species along with about the frequency of cross-species transmission events (Morga-Jacquot et al., 2021).

In the present study, high-throughput deep sequencing was used to characterize the genetic diversity of OsHV-1 found in both *O. edulis* and *M. gigas*. To this end, forty new OsHV-1 were sequenced and assembled. Using these data and previously gathered information about lineages circulating in different host species, our aims were to (*i*) describe OsHV-1 genomic diversity patterns in both species, (*ii*) estimate the genetic isolation between lineages infecting both species and (*iii*) determine if OsHV-1 cross-species transmission occurred and at which frequency. To do so, a combination of comparative and population genomic, phylogenetic and phylodynamic approaches were applied and results are discussed in the context of current knowledge of OsHV-1 molecular epidemiology.

## 2. Materials and Methods

### 2.1. OsHV-1 sampling

To study the diversity of OsHV-1 in different host species, archived samples from Ifremer National Reference Laboratory and European Union Reference Laboratory for mollusk diseases collection were used. OsHV-1 viruses were sampled from Pacific oysters *M. gigas* (seventeen samples) collected in the field during OsHV-1 infection outbreaks between 2005 and 2010 from individuals of different developmental stages (larvae, spats and adults) at five sites along the French coast (Antioche inlet, Palavas lagoon, Thau lagoon, Brittany inlet, Pirou Agon-Coutainville). Wild flat oysters *O. edulis* (twenty-three samples) collected in the field from April to August between 2007 and 2009 in the Quiberon bay as describe by (Arzul et al., 2011) and one sample has been collected in Sweden (Table S1, Fig. 1A and 1B). This last sample was used to compare the viral lineage from France and Sweden within *O. edulis* individuals to widen the geographical area studied. Criterion used to select *M. gigas* samples was *i)* the geographic proximity to *O. edulis* samples and *ii)* the year of samples collection, which covers the *O. edulis* samples collection period. Before DNA extraction, animals have been stored at -80°C or in ethanol from 6 to 14 years.

**Figure 1.**
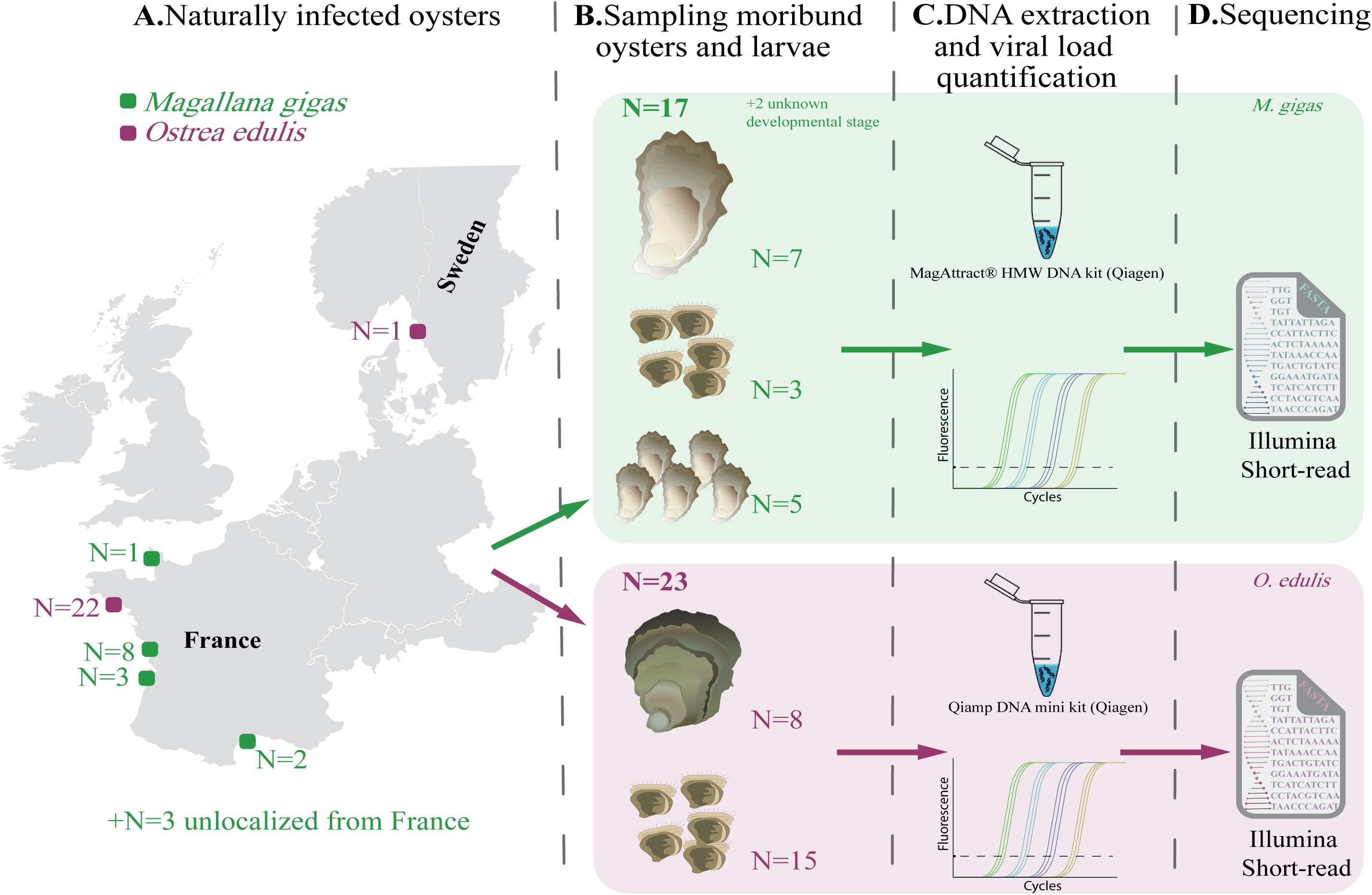
Schematic overview of the experimental sampling design. (A-B) Larvae, spat, and adults naturally infected with the virus were collected from *M. gigas* and *O. edulis* in France and Sweden. (C) DNA was extracted from individual tissues and the viral load was assessed by qPCR. D) DNA samples were sequenced using the Illumina short-read technology. *M. gigas* and *O. edulis* samples and treatments are represented in green and purple, respectively.

Samples were named according to the previously recommended nomenclature (Kuny & Szpara, 2022) which is made of the virus name, the sampling location, the date of collection, the host species, and the sample number (*e.g.* OsHV-1/Rivedoux.FRA/2008/Cg-S-053 for the sample number 053 collected in 2008, in Rivedoux, from *M. gigas* spat). However, hereafter, shortened names have been used to facilitate reading (*e.g*. CG-08-Ri-S-053 for the same sample, see Table S1 for correspondence).

### 2.2. DNA extraction, viral copy numbers quantification and sequencing

Depending on the host species, two different methods were used for samples DNA extraction (Fig. 1C). For *M. gigas*, DNA was extracted from mantle of moribund individual spats, from pool of larvae or from pool of shred oysters using the Qiamp DNA Mini kit (Qiagen) according to the manufacturer’s protocol. For *O. edulis,* DNA was extracted from gills of adult individuals or from pool of larvae using the MagAttract^®^ HMW DNA kit (Qiagen) according to the manufacturer’s protocol.

DNA purity and concentration were assessed using a Nano-Drop ND-1000 spectrometer (Thermo Scientific) and Qubit^®^ dsDNA BR assay kits (Molecular Probes Life Technologies), respectively. Then quantification of viral copy numbers was carried out in duplicate by quantitative PCR using a Mx3005 P Thermocycler (Agilent) as described by (Pepin et al., 2008). Each reaction contained 1µl of Milli-Q Water, 2µl of each primer at 5,5 µM, OsHVDP For (forward) 5′-ATTGATGATGTGGATAATCTGTG-3′ and OsHVDP Rev (reverse) 5′-GGTAAATACCATTGGTCTTGTTCC-3′, 10µl of Brilliant III Ultra-Fast SYBR® Green PCR Master Mix (Agilent), and finally 25 ng of DNA sample from infected oysters or Milli-Q water (non-template control) in a total volume of 20µl. The qPCR program consisted in 3 min at 95 °C followed by 40 cycles of amplification at 95 °C for 5s and 60 °C for 20s. Results were expressed as the log-transformed copy number of viral DNA per nanogram of extracted DNA (cp.ng^-1^).

Illumina DNA-Seq library preparation and sequencing were performed at four sequencing platforms (Fig. 1C, see Table S1 for details about libraries preparation and sequencing technologies). Libraries were prepared using (*i)* True seq DNA PCR free kit, (*ii)* Shotgun PCR-free library preparation kit (Lucigen) or (*iii*) Illumina genomic PCR-free kit. The sequencing step was performed using (*i*) HiSeq^TM^ 2500 device (paired-ends, 100 pb) by GenoToul (Toulouse, France), (*ii*) HiSeq^TM^ 4000 device (paired-ends, 150 bp) by the Lille Integrated Genomics Advanced Network for personalized medicine (LIGAN-PM) sequencing platform (UMR 8199, CNRS, Lille, France) or (*iii)* NovaSeq^TM^ 6000 device (paired-ends, 150 bp) by the Genome Quebec Company (Genome Quebec Innovation Center, McGill University, Montreal, Canada) or by Fasteris Life Science Genesupport (Plan-les-Ouates, Switzerland).

### 2.3. De novo assembly of OsHV-1 genomes

*De novo* OsHV-1 genome assemblies were obtained by processing sequencing reads as previously described (37,38 and see 48 for details on the bioinformatic pipeline). Briefly, for each library, raw reads were filtered according to their quality (PHRED score > 31) and adapters trimmed using Fastp v0.20.1 (with parameter -q 31; (Chen et al., 2018). In order to estimate the number of OsHV-1 reads and coverage in each sample, filtered reads were aligned to the reference genome of OsHV-1 (accession number: NC_005881.2; (Davison et al., 2005) using Bowtie2 v2.4.1 (parameter --local and --no-unal; 50). Meanwhile, reads were aligned to the reference genome of *M. gigas* (accession number: GCF_902806645.1, (Peñaloza et al., 2021)) or *O. edulis* (accession number: [GCA_024362755.1](https://www.ncbi.nlm.nih.gov/datasets/genome/GCA_024362745.1/), (Boutet et al., n.d.)) according to the species from which the sample was collected, using Bowtie 2 v2.4.1 (parameter --local). Non-oyster reads were selected with SAMtools v1.9 (option view -f 4, (Li et al., 2009)) and Seqtk v1.2 subseq (default parameters; (Li, 2013)). Non-oyster reads were then *de novo* assembled with metaSPAdes v3.15.4 (parameter --meta; (Bankevich et al., 2012)). OsHV-1 contigs were selected using BLASTn v2.12.0 (options -max_target_seqs 1, -evalue "1e-10"; (Altschul et al., 1990)) against the OsHV-1 reference genome. OsHV-1 contigs were then manually re-ordered in Geneious Prime v2022.2.2 by visualizing reads alignment on the *de novo* assembled contigs to identify overlaps between trimmed reads at each contig extremities. To avoid inverted repeat regions reconstruction issues for all subsequent analyses, every OsHV-1 genome was reconstructed as non-redundant (NR) genome containing only one copy of each duplicated regions (IR_L_ and IR_S_) as previously described (Delmotte-Pelletier et al., 2022; Dotto-Maurel-Pelletier et al., 2022; Morse et al., 2017). NR-genomes were validated and corrected by similarity matrix visualization of each genome to itself using the dotplot tool (based on the EMBOSS v6.5.7 tool dottup) implemented in Geneious Prime v2022.2.2. To verify and clean the final assemblies as well as to assess coverage uniformity, quality filtered reads were aligned back to the new constructed OsHV-1 NR-genomes using Bowtie 2 v2.4.1 (parameter --end-to-end).

### 2.4. Genomic architecture and ORF predictions

The genomic annotation in this study has been carried out at genome and loci scales, including the identification of genomic regions (U_L_, IR_L_, X, IR_S_, U_S_), ORF prediction and functional annotation, signal peptides and transmembrane domain prediction.

At the genome scale, the structural genomic regions annotation, was performed using a combination of tools. Firstly, BLASTn v2.12.0 (Altschul et al., 1990) implemented in Geneious Prime v2022.2.2 was used to compare each OsHV-1 NR-genome assembled in this study with the OsHV-1 reference genome (NC_005881.2) and transfer raw annotation to assembled NR-genome. Secondly, the structural annotations were corrected using coverage plots generated after remapping reads to control and clean the assembly. By observing where the coverage levels experienced changes or abrupt shifts, it allows the identification and correction of the exact boundaries between repeated and unique regions.

#### 2.4.1. Ancestral sequence reconstruction

To ensure consistency in the predictions across different genomes and to facilitate the identification of polymorphic sites and gene positions, an ancestral sequence of OsHV-1 for each host species was reconstructed. To this end, first a multiple alignment of genomes sampled from *M. gigas* (twenty genomes) or from *O. edulis* (twenty-three genomes) was obtained using MAFFT v1.4.0 (Katoh et al., 2002). Subsequent analyses were performed using the model of nucleotide substitution HKY+G+I based on Akaike Information Criterion (AIC) obtained with jModelTest (Darriba et al., 2012) implemented in MEGA v11.0.11 (Tamura et al., 2021). Ancestral sequences of studied OsHV-1 genomes from both host species were reconstructed using Beast v1.10.4 (Drummond & Rambaut, 2007). Different combinations of coalescent models (*i.e.* constant size, GMRF Bayesian skyride, expansion growth, and exponential growth) and molecular clocks (*i.e.* strict clock and uncorrelated relaxed clock) were tested to find out the most appropriate model. The maximum likelihood estimator calculated by path sampling and stepping-stone sampling were compared to select the most appropriate model. According to results obtained, ancestral sequences were reconstructed using a strict clock and a constant size coalescent model for the viruses from *M. gigas*, and an uncorrelated relaxed clock and an expansion growth model for the viruses from *O. edulis* and MCMC chains of 10,000,000 steps were ran and sub-sampled every 1,000,000 generations.

#### 2.4.2. ORF, signal peptides and transmembrane domains predictions

Subsequently, local predictions of genes, signal peptides and transmembrane domain on the two reconstructed ancestral sequences were performed. ORFs were predicted with Prodigal v2.6.3 (Hyatt et al., 2010) with default parameters, a tool already used for double-stranded DNA viruses infecting eukaryotes (González-Tortuero et al., 2021). Functional annotations of predicted proteins were performed with EggNOG-mapper v2.1.8 (Cantalapiedra et al., 2021). Signal peptides were predicted using SignalP v6.0 (Teufel et al., 2022) and transmembrane domains using DeepTMHMM v1.0.24 (Hallgren et al., 2022).

### 2.5. Within samples genomic diversity

#### 2.5.1. Analysis of intra-sample single nucleotide variants rarefaction

Since some samples, particularly those containing larvae, include multiple individuals, we will refer to intra-sample diversity rather than intra-individual diversity. To verify that sequencing errors was not a major driver of intra-sample diversity and investigate the degree to which sequencing effort affected the number of intra-sample variations detected, a rarefaction analysis on the number of detected variations was performed for each sequencing library (Cameron et al., 2021; Delmotte-Pelletier et al., 2022). To this end, an iterative variant calling analysis was performed on subsampled reads that were mapped on the previously assembled genomes. In order to accelerate analyses, a nonlinear subsampling incrementation was performed based on 1%, 2%, 3%, 4%, 5%, 6%, 7%, 8%, 9%, 10%, 15%, 20%, 25%, 30%, 50%, 80% and 100% of the total number of reads of libraries. After each subsampling/variant calling step, variants were classified as intra-sample Single Nucleotide Variants (iSNVs) and insertion or deletions (InDels) and counted in order to increment rarefaction curves.

#### 2.5.2. Variant calling

Then, to determine the genomic diversity within samples, variations were called against each of the 40 genomes previously assembled, using Freebayes v1.3.5, (Garrison & Marth, 2012) with the following settings: --use-mapping-quality, --min-repeat-entropy 1, --haplotype-length 0, --min-alternate-count 5,--pooled-continuous, --hwe-priors-off, --allele-balance-priors-off and --gvcf. All the variations were then normalized, multiallelic variants and biallelic blocks were decomposed and then sorted and deduplicates (using programs normalize, decompose, decompose_blocksub, sort and uniq) with vt v2015.11.10 (Tan et al., 2015) using default parameters. iSNVs detected by Freebayes were mapped on the corresponding consensus, then all the forty-one consensuses with their corresponding annotated iSNVs were aligned using MAFFT v1.4.0 (Katoh et al., 2002) implemented in Geneious Prime to re-coordinate all iSNVs. Re-coordinated iSNVs were then used to create unique identifier as described by Delmotte-Pelletier et al. (2022). Briefly, unique identifiers are composed of *(i*) the position of the iSNVs along the multiple alignment, (*ii*) the reference allele (REF) and *(iii*) the alternative allele (ALT) (*i.e.* position_REF >ALT).

#### 2.5.3. Statistical description of the intra-sample diversity

Multi-Dimensional Scaling (MDS) plots were used to examine the impact of host species on the intra-sample genetic diversity of OsHV-1. The first MDS analysis summarized the allelic frequencies of each allele within the sequencing libraries, allowing to explore the distances between pairs of iSNVs in terms of their allelic frequencies within each library. This information was linked to those of host species for the visualization step. The second MDS was performed using a spearman correlation matrix based on individuals’ genetic similarities. The number of clusters of samples were searched using the kmeans method and were represented on the MDS plot. Then, an allelic association analysis was performed using the co-occurrence matrix of alleles among samples, as well as the adjacency and incidence of vertices on each other. Connections between edges and their weights were used to represent the strength of interaction between individuals in the MDS graph.

Both MDS and plotting were performed in R v4.2.2 using stats v4.2.2 (R Development Core Team, 2005), igraph v1.3.5 (Csardi & Nepusz, 2006) and ggplot2 v3.4.0 (Wickham, 2009) packages. To examine the density of iSNVs within each genomic region (*i.e*. U_L_, IR_L_, X, IR_S_, U_S_), the ratio of the number of iSNVs detected within a region to the length of the region 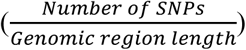 has been calculated.

### 2.6. Whole genome alignment and recombination detection

#### 2.6.1. Identification and description of OsHV-1 structural variations

To compare the genomic architecture between viral lineages and determine large-scale evolutionary events, the forty reconstructed NR-genomes were aligned with three published OsHV-1 genomes (OsHV-1 µVarA, *M. gigas,* 2010, France: KY242785, OsHV-1 µVarB, *M. gigas,* 2011, Ireland: KY271630, and OsHV-1 Ref., *M. gigas*, 1995, France: NC_005881.2) using Mauve v1.1.3 (progressive Mauve algorithm, with default parameters, (Darling et al., 2004)). These three genomes were available in public databases and were also assembled from Pacific oysters. Furthermore, the lineages are very similar to those detected in our libraries. However, the three published genomes assembled from clams or scallops were not include as they are too distant from the lineages studied in this research, and not enough genomes are available for these species. Our multiple alignment of the forty-three OsHV-1 genomes allowed us to identify typical genomic structures based on rearranged blocks and to classify genomes according to their structure. Specifically, five distinct genomic structures were identified in this study.

To plot the five types of genomic structures observed in our samples, pairwise alignments of the five representative lineages against the OsHV-1 reference genome (NC_005881.2) were conducted using minimap2 v2.24 (parameters -ax asm5 --eqx; (Li, 2018)). Alignments were then sorted and indexed using SAMtools v1.14 (Li et al., 2009). Structural rearrangements were characterized using Syri v1.5.4 (Goel et al., 2019) and plot with the python package plotsr v0.5.3 (Goel & Schneeberger, 2022).

#### 2.6.2. Manual restructuring of OsHV-1 genomes and multiple alignment

As some genomes contained large rearrangements, thirty assembled genomes were manually modified, from the Mauve multiple alignment, in Geneious Prime to ensure that the genomic architecture was identical for every genome, making them comparable to each other. Genomes were then aligned using MAFFT v1.4.0 (Katoh et al., 2002) that allowed the computation of nucleotide identity percentages between pair of genomes.

#### 2.6.3. Recombination events detection

Gubbins v3.3 (Croucher et al., 2015) with default settings was used to identify regions that might be impacted by recombination events among the assembled OsHV-1 NR-genomes. Additionally, to confirm results obtained with Gubbins, linkage disequilibrium values (D’) were computed to test for random association of allelic distribution and then explore the possibility of recombination in our dataset. D’ values were computed in DNAsp6 (Rozas et al., 2017) based on informative sites and significance was assessed using a Fisher’s exact test. D’ values were then plotted as a heatmap in R using the heatmap function from the ggplot package v3.4.0 (Wickham, 2009). All genomic regions detected as potentially recombinant were removed from the downstream analyses (Table S5).

### 2.7. Consensus based diversity computation

#### 2.7.1. Nucleotide diversity and segregating sites

Standard indices of diversity Θ_π_ (*i.e.* measure of the average number of nucleotide differences between pairs of DNA sequences) and S (*i.e.* the number of polymorphic/segregating sites along the alignment of sequences) were computed to estimate the genetic diversity among all samples, among *O. edulis* samples and among *M. gigas* samples. Diversity indices Θ_π_ and S were computed in R using the functions nuc.div and seg.sites from the pegas package v1.3 respectively (Paradis, 2010).

#### 2.7.2. Population structure and phylogeny

First, a phylogenetic analysis has been performed to explore the population structure and evolutionary relationships among OsHV-1 lineages. A phylogenetic search that applied a maximum-likelihood approach was performed on the alignment of the forty-three genomes using PhyML v3.3.2 (Guindon et al., 2010) with 1000 bootstraps replications. The best fitting model of evolution for the samples was found using jModelTest (Darriba et al., 2012) in MEGA v11.0.11 (Tamura et al., 2021) based on AIC. Therefore, a HKY+G+I model was used. Results were visualized using Fig.Tree v1.4.4 (available at http://tree.bio.ed.ac.uk/software/Fig.tree/).

#### 2.7.3. Investigating OsHV-1 genetic differentiation between host species

Description of consensus-based diversity was done using population genetic statistics such as the deviation of allelic frequencies from the neutrality equilibrium (Tajima’s D) and the differentiation level of viruses infecting the two host populations (G_ST_).

To obtain insights into deviation from demographic equilibrium and/or selective neutrality, Tajima’s D values were computed in DNAsp6 v6.12.03 (Rozas et al., 2017) for 1kb windows along the alignment of NR-genomes, in order to standardize the amount of available information among analyses. Tajima’s D values were calculated for the overall population (forty-three genomes) as well as for samples from each host species separately. Under the neutral evolutionary model, Tajima’s D is expected to be null. At a locus scale, positive Tajima’s D values can be induced by balancing selection, due to the maintenance of highly divergent variants and, negative Tajima’s D values can be the result of a bottleneck or of a recent selective sweep, which both result in an excess of weakly divergent alleles. At the whole genome scale, positive Tajima’s D values reflect a decrease of the population effective size thus favoring intermediate frequency alleles, while negative Tajima’s D values suggest an increase of the population effective size implying expansion of rare alleles.

Then, differentiation measures were computed between viral populations from both host species as they reflect the genetic diversity among each species relative to the diversity of the whole sampling. The pairwise Nei G_ST_ method was used and, per locus G_ST_ values were computed for 1kb windows with the option diff_stats in R using the package mmod v1.3.3 (Winter, 2012). G_ST_ values were also computed for 100bp windows within 1kb windows that were detected as differentiated to define precisely regions involved in differentiation and facilitated the link with ORF potential function for biological hypothesis assessment.

An AMOVA was conducted to investigate the possible factors contributing to the structuring of OsHV-1 genomic diversity at the consensus level. To perform the AMOVA, the global alignment of NR-genomes was first converted into a distance matrix using the Tamura-Nei model implemented in the dist.multidna function of the R package ape v5.8-1 (Paradis et al., 2024). The analysis was then conducted using the poppr.amova function from the poppr package v2.9.6 (Kamvar et al., 2014) in R. Grouping factors were defined based on metadata associated with each NR-genome, such as geographic origin, year of sampling, and host species. The statistical significance of the observed variance components was tested using 1,000 permutations. This approach allowed us to quantify the proportion of genetic variation explained by each factor and to identify which ones significantly contributed to the observed structuring of OsHV-1 genomic diversity.

#### 2.7.4. Detection of selection signatures

To explore the potential effects of natural selection, signals of selective sweeps were searched in our dataset. A selective sweep occurs when a new beneficial mutation increases in frequency and becomes fixed, resulting in a reduction or elimination of genetic variation near the mutation. RAiSD v2.9 (Alachiotis & Pavlidis, 2018) was used to detect selective sweep signatures along OsHV-1 genomes. The RAiSD tool utilizes the µ-statistic, a test that scores genomic regions by measuring changes in the Site Frequency Spectrum (SFS), levels of Linkage Disequilibrium (LD), and the amount of genetic diversity along a genome (Alachiotis & Pavlidis, 2018).

#### 2.7.5. Ancestral state reconstruction of the host species trait

In order to analyze the temporal signal and the ‘clocklikeness’ of the maximum likelihood tree and first explore the evolutionary rate of OsHV-1, TempEst v1.5.3 (Rambaut et al., 2016) has been used to plot the divergence of each tip from the root over time. The root-to-tip divergence (RTTD) plot has been performed separately for *O. edulis* viruses (twenty-three genomes) and for the two subclusters of the OsHV-1 *M. gigas* viruses (respectively nine and eleven genomes) using a midpoint rooted tree and a best fitting root approaches for comparison.

Analyses were performed using a HKY+G+I model of nucleotide substitution based on AIC obtained using jModelTest (Darriba et al., 2012) in MEGA v11.0.11 (Tamura et al., 2021). As before, different combinations of coalescent models (*i.e.* constant size, GMRF Bayesian skyride, and exponential growth) and molecular clocks (*i.e.* strict clock and uncorrelated relaxed clock) were tested to find the most appropriate model for our dataset. The maximum likelihood estimator calculated by path sampling and stepping-stone sampling for each model were compared. According to those, an uncorrelated relaxed molecular clock and a constant size coalescent model has been and a MCMC chain of 10,000,000 steps was run and sub-sampled every 1,000,000 generations using Beast v1.10.4 (Drummond & Rambaut, 2007). The effective sample size, representing the mixing of the population (Ne > 200), and the stationary distribution of the trace were checked in Tracer v1.7.2 (Rambaut et al., 2018) for all parameters, using an initial burn-in of 10%. Finally, maximum clade credibility (MCC) trees were generated from tree output files from BEAST in TreeAnnotator v1.10.4 (Drummond & Rambaut, 2007), and annotated in Fig.Tree v1.4 (http://tree.bio.ed.ac.uk/software/Fig.tree/).

#### 2.7.6. Modeling of demographic evolution

To assess the historical population dynamics of OsHV-1 in both host species, a demographic analysis was conducted by reconstructing past changes in population size over time. To do so, the log files of the trees that were previously generated to reconstruct ancestral sequences of OsHV-1 in both host species were used. The demographic histories were then reconstructed using a GMRF Skyride model in Tracer v1.7.2 (Rambaut et al., 2018).

## 3. Results

### 3.1. Detection of OsHV-1 in O. edulis and M. gigas collection samples

The viral genome copy number varied from 1.47×10^3^ to 3.07×10^8^ copies per ng of DNA, with an average quantity of 1.34×10^7^ copies per ng of DNA for the samples collected from *O. edulis* (Fig. S1, Table S1). For samples collected from *M. gigas*, the viral genome copy numbers ranged from 2.21×10^4^ to 4.21×10^7^ copies per ng of DNA, with an average of 2.81×10^6^ copies per ng of DNA (Fig. S1, Table S1).

As the minimum threshold for efficient Illumina sequencing of virus particles is 10^3^ copies, all samples were found to be suitable for sequencing (Delmotte-Pelletier et al., 2022; Dotto-Maurel-Pelletier et al., 2022; Morga-Jacquot et al., 2021).

### 3.2. Sequencing statistics

Sequencing depth and viral genome coverage were both correlated to the total number of reads for the different libraries and to the OsHV-1 genomes copy number respectively (Fig. S1). Illumina sequencing step produced from 18.9 to 290.9 million reads for a given sample (average 112.2 M reads ± 60.3 SD, Fig. S1 and Table S1). Host reads represented from 5.2% to 91.2% of the total reads. The percentage of reads mapped to OsHV-1 varied from 0.013% to 38.51%, and the genome coverage varied accordingly from 12 to 26,790 reads depth. Using this sequencing dataset, 40 OsHV-1 Non-Redundant genomes (NR-genomes) were *de novo* assembled. The length of the assembled NR-genomes varied from 185.9 kbp to 190.2 kbp.

To verify that sequencing errors were not a major driver of detected diversity and to investigate the degree to which sequencing effort affected the number of intra-sample variations detected, a rarefaction analysis on the number intra-sample single nucleotide variants (iSNVs) was performed. Rarefaction curves reached a plateau at 150,000 reads corresponding to an average read depth of 110 (Fig. S2). Therefore, this value was used as the minimum number of reads necessary to accurately detect nucleotide polymorphisms in our samples. 12 libraries did not reach this threshold and were excluded from all analyses focusing on intra-sample diversity. For other libraries, when a high number of sequences were used, no additional variations were detected, which would have been the case if there was an accumulation of sequencing errors.

### 3.3. Intra-sample OsHV-1 diversity

Based on variant calling results, a total of 776 iSNVs - defined as sites displaying at least two alleles with frequencies greater than 10% in at least one sample - were detected across genomes. Among these, 648 sites exhibited two alleles, 85 sites exhibited three alleles, 25 displayed four, and 18 showed more than four alleles (gaps) (Fig. 2A, Table S2). Similarly, across OsHV-1 genomes obtained from *O. edulis* samples, 496 iSNVs were identified, 69 of which presenting more than two alleles. On the other hand, OsHV-1 genomes obtained from *M. gigas* samples harbored 329 iSNVs, with 53 of them carrying more than two alleles. A total of 51 sites were detected as iSNVs in both species.

**Figure 2.**
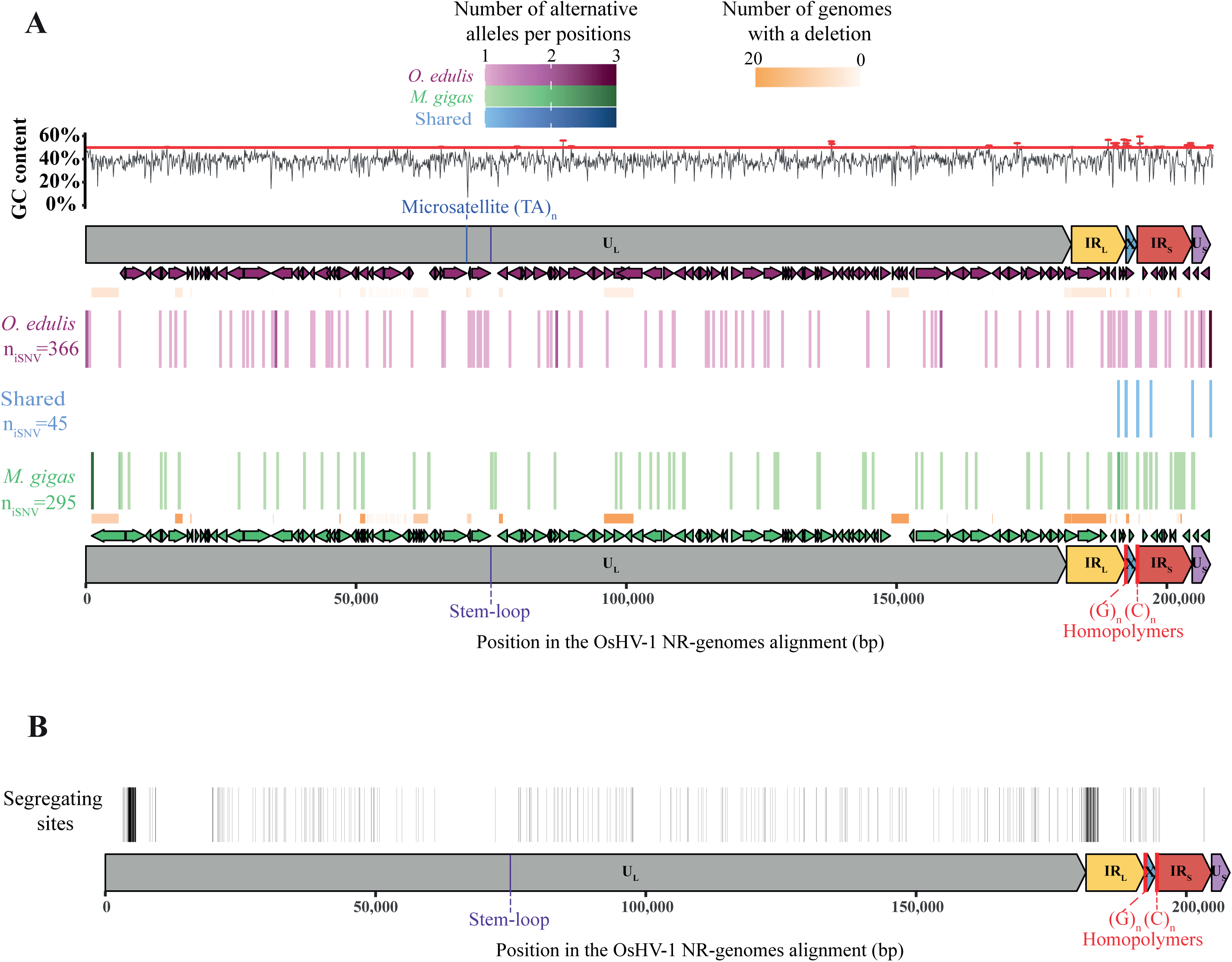
Distribution of iSNVs and segregating sites across the OsHV-1 genome. (A) Distribution of intra-sample single nucleotide variants (iSNVs) along the alignment of OsHV-1 *de novo* assembled genomes. iSNVs detected exclusively in the *M. gigas* population are shown in green, those detected in the *O. edulis* population in purple and iSNVs shared by both populations in blue. The intensity of the point reflects the number of alleles detected at the corresponding genomic position. Positions of iSNVs are plotted relative to annotated genomic regions. GC content is represented in grey, with values exceeding 50% highlighted in red. (B) Distribution of segregating sites along the alignment of OsHV-1 *de novo* assembled genomes.

Out of the 496 iSNVs detected in *O. edulis* samples, 336 were located within 58 predicted ORFs (Table S3). More precisely, 161 iSNVs were found in ORFs coding for proteins with unknown functions, while the remaining iSNVs were identified in ORFs coding for proteins with known domains or functions (Table S3). In detail, 105 iSNVs were detected in ORF coding for putative transmembrane proteins (in ORFs 22, 41, 54, 57, 59, 65, 72, 77, and 111), 26 in ORF coding for putative DNA primase (ORFs 66 and 7), 15 in ORF coding for putative DNA packaging protein (ORF 109), 6 in ORF coding for DNA polymerase (in ORF 100), and 6 in ORFs coding for Zinc-finger RING type protein (ORFs 9, 38, 97, and 117).

In viral lineages sampled from *M. gigas*, 203 out of the 496 detected iSNVs were located within 44 predicted ORFs. 57 iSNVs were found in 21 ORFs coding for proteins with unknown functions, while the remaining 148 iSNVs were identified in ORFs coding for proteins with known domains or functions. More precisely, 81 iSNVs were detected in ORFs coding for transmembrane proteins (ORF 4, ORF 22, 41, 65, 68, and 77), 48 were detected in ORFs coding for membrane proteins (ORF 16, 36, and 63) and 6 in ORFs coding for Zinc-finger, RING type. The remaining iSNVs were detected in ORFs coding for DNA primase, DNA packaging, or DNA polymerase (Table S3).

In proportion to the size of the regions, the U_L_ region exhibits fewer iSNVs (0.14% for *M. gigas* samples and 0.19% for *O. edulis* samples) compared to the other regions (Table S4). In the *M. gigas* samples, the genomic regions with the highest iSNV density are the IR_L_ and IR_S_ regions, containing 0.81% and 0.57% of the iSNVs, respectively, while 0.53% of the X and U_S_ regions contain iSNVs. In the *O. edulis* samples, the OsHV-1 U_S_, IR_L_, and X genomic regions exhibit the highest iSNV density, containing 1.03%, 0.99%, and 0.72% of the iSNVs, respectively, while only 0.36% of the IR_S_ region contains iSNVs.

Interestingly, the iSNVs shared by samples from both oyster species are located in the repeated regions IR_L_ and IR_S_, particularly in regions that exhibit a GC content greater than 50% (Fig. 2B). No shared iSNVs is detected in the X region, except in the homopolymers at the junctions with the repeated regions, which display an accumulation of iSNVs.

The multidimensional scaling (MDS) analysis of allelic frequencies of polymorphic sites showed that alleles frequencies in samples from *O. edulis* are different from those in samples from *M. gigas*. The MDS further indicate a clear structuring of intra-sample variations according to host species (Fig. 3A). However, 92 iSNVs were more similar within both species compared to others, indicating that some positions are variable independently of the species studied (Fig. 3A).

**Figure 3.**
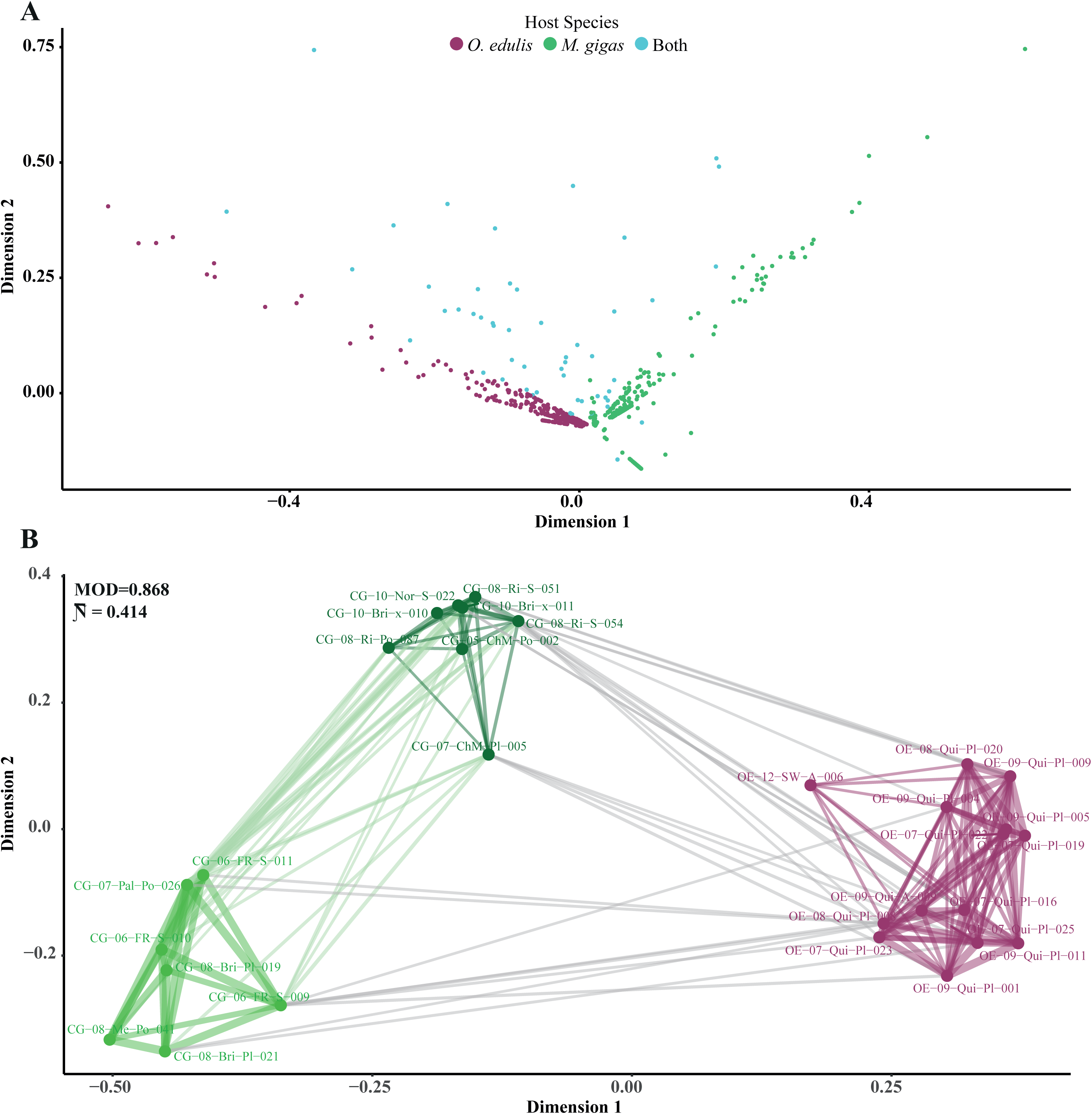
From intra sample population diversity to host specialization. (A) Multidimensional scaling (MDS) plot showing the distribution of allelic frequencies at polymorphic sites. Each point represents an allele, colored according to the host species in which it was detected (*O. edulis* in purple, *M. gigas* in green). (B) MDS plot representing the similarity between OsHV-1 lineages within oysters according to the allelic frequencies of the iSNVs they harbor. This two-dimensional projection provides a visual representation of the genetic structuring of viral populations across hosts. Samples from *O. edulis* are shown in purple, while those from *M. gigas* are shown in green and form two distinct clusters. Lines connecting samples represent the degree of similarity between their viral populations, and its color indicates whether individuals belong to the same host species.

The allelic association graph, coupled with the MDS analysis, summarized the two-dimensional comparison of the co-occurrence frequency of each iSNVs among samples (Fig. 3B). The modularity values of a network indicate the strength of connections within and between communities, and thus, provide information about the quality of network division. The analysis of alleles co-occurrence revealed the presence of three distinct clusters based on modularity measurements. Out of these three clusters, two corresponded to *M. gigas* samples (represented by the dark and light green clusters on Fig. 3B) and one cluster corresponding to *O. edulis* samples (represented by the purple cluster on Fig. 3B). The dark green cluster of *M. gigas* samples comprises samples collected from various developmental stages between 2005 and 2010 in Charente-Maritime, the Breton inlet, and Normandie. On the other hand, the light green cluster of *M. gigas* samples consists of samples collected between 2006 and 2008 in Palavas and Thau lagoons, and the Breton inlet, encompassing all developmental stages.

### 3.4. Structural variations

Using a whole genome alignment approach, a structural analysis of all OsHV-1 NR-genomes assembled from *M. gigas* and *O. edulis* samples have been performed (Fig. S3). The whole genome comparison revealed six types of genomic architecture (Fig. 4). The first one includes four NR-genomes obtained from *M. gigas* and that are similar to the reference genome described by (Davison et al., 2005), characterized by a size of approximately 190 kbp. The five other structural types were identified and described in comparison with the OsHV-1 reference genome (Davison et al., 2005). First, seven NR-genomes of 190 kbp collected from *M. gigas* structurally differ from the OsHV-1 reference genome by a reversion of 8,530 bp in the U_L_ region between ORF 36 which is disrupted (potentially not functional) and ORF 43. Second, nine NR-genomes of 204 kbp also sampled from *M. gigas* were analogous to the µVar genotype (Burioli et al., 2017) which contains four deletions in the U_L_ region (ranging from 606 bp to 3,550 bp), one in the IR_L_ (725 bp) region, and one insertion of 2,671 bp in the U_L_ region between ORF 43 and 44, for a total of approximately 10 kbp of insertions or deletions (InDels). Third, one NR-genome of 192 kbp, sampled from *O. edulis,* contains a reverse translocation of the first part (between ORF 6 and ORF 4 which is disrupted) of the TR_L_ region (7 kbp) and an insertion of 2,204 bp upstream to the U_L_ region. In addition, this genome contains multiple large deletions (6 deletions for a total of 5,303 bp) and insertions (5 insertions for a total of 4,017 bp) for an average of 13 kbp InDels. Fourth, twenty NR-genomes of 185 kbp and sampled from *O. edulis*, contain a reverse translocation of the first part of the TR_L_ region (4 kbp). In addition, these genomes contain multiple large deletions (11 deletions for a total of 6,423 bp) and insertions (4 insertions for a total of 1,133 bp) for an approximate 11 kbp of InDels in total. Finally, two NR-genomes contain the same reverse translocation, deletions, and insertions as those detected in the fourth structural type with an additional inversion of 1,470 bp was detected in the UL region (ORF 50 disrupted) for a total of approximately 12 kbp of variations.

**Figure 4.**
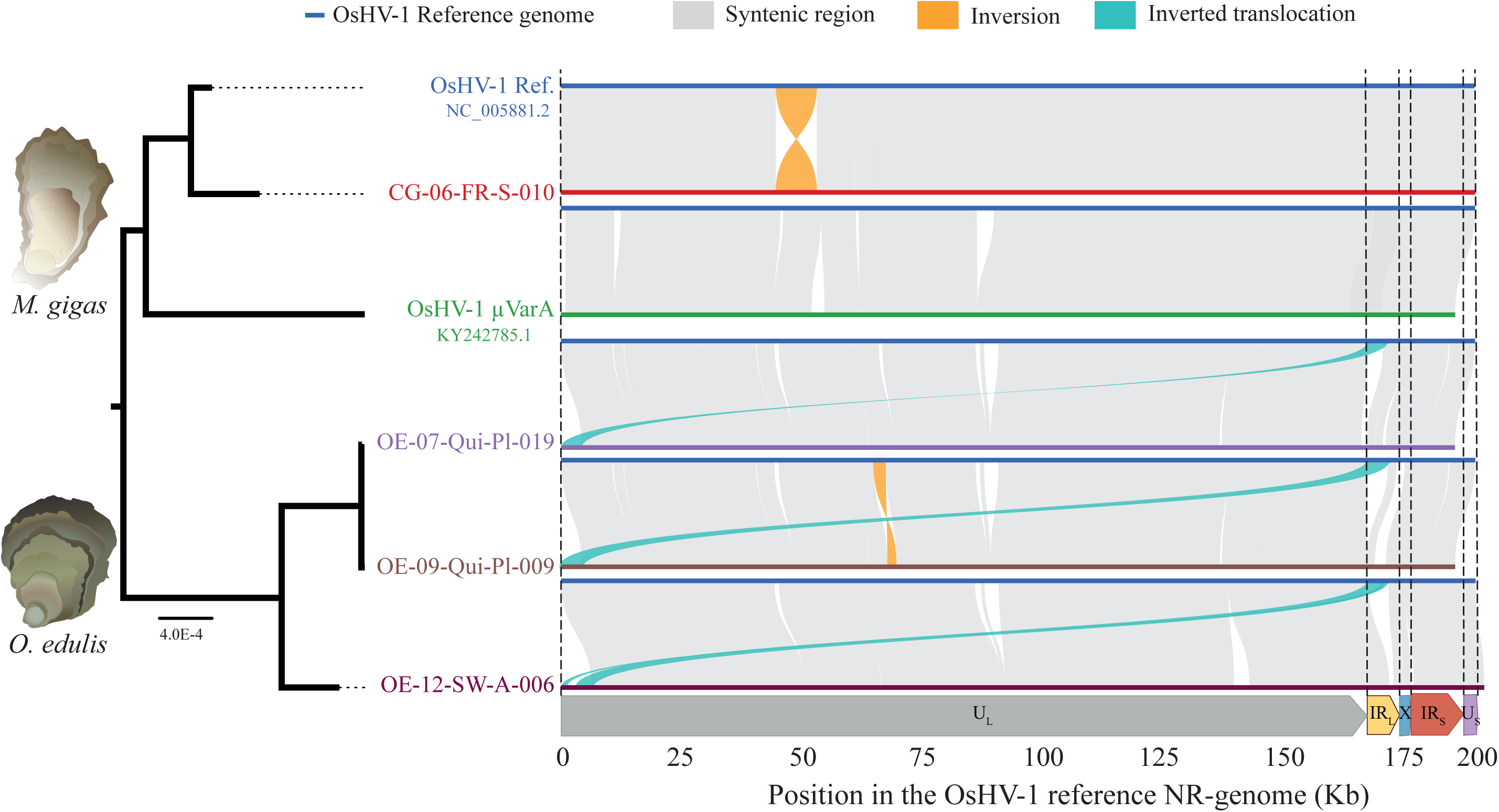
Structural variations among OsHV-1 genomes. Structural genome comparisons of six representative OsHV-1 genomes are shown alongside their phylogenetic relationships, inferred using a maximum likelihood approach (Guindon et al., 2010). The description of genomic structural variations was based on a pairwise comparison to the OsHV-1 reference genome (NC_005881.2, Davison et al., 2005). In each comparison, the reference genome is represented by a dark blue bar. Conserved syntenic regions are shown in light grey, inversions in orange, and inverted translocations in light blue. The lower panel illustrates the genomic architecture and positional context of the structural variants along the OsHV-1 reference NR-genome.

### 3.5. Recombination hotspots and alignment curation

To ensure that homologous regions are all in phase, and that multiple alignment and phylogenetic analyses can be carried out, genomic regions of the forty-three OsHV-1 NR-genomes have been manually rearranged. The recombination analysis identified the occurrence of 5 potential recombination events along OsHV-1 genomes, that may have impacted two genomic regions (Fig. S5). The first potential recombination event was located in the U_L_ region between 61,474 pb and 68,405 bp, specifically between a microsatellite (TA)_n_ present in the viral genomes collected from *O. edulis* and a stem loop predicted to be the probable origin of replication of OsHV-1 (Davison et al., 2005). The other four recombination events detected were located in repeated regions detected respectively between 173,860 and 175,357 bp at the junction between U_L_ and IR_L_ regions, between 178,966 and 180,440 bp and from 184,844 to 185,190 bp both in the IR_L_ region, and finally from 194,785 to 195,422 bp located in the IR_S_ region. In complement, D’² values were mostly close to 1, indicating strong linkage disequilibrium (Fig. 5) and suggesting low recombination rates. However, one position close to the 3’ end of the genome within the ORF 6 (position 3,108), as well as two other positions at the 5’ end of the U_L_ respectively within ORF 88 and ORF 104 (positions 137,500 and 164,760), one position in the IR_L_ within ORF 114 (position 182,122), and one position in the IR_S_ (position 195,034) regions had very low linkage disequilibrium values, close to 0. None of the detected low D’² values was in a homopolymer region but two low D’² positions (positions 182,122 and 195,034) correspond to recombination breakpoints detected by Gubbins (Table S5). As a consequence, a total of 11,184 bp distributed within 13 regions potentially involved in recombination were removed from the alignment for the subsequent analyses, this correspond to 541 SNPs that have been removed from consensus-scale analyses. The suppressed regions are referenced in the Table S5.

**Figure 5.**
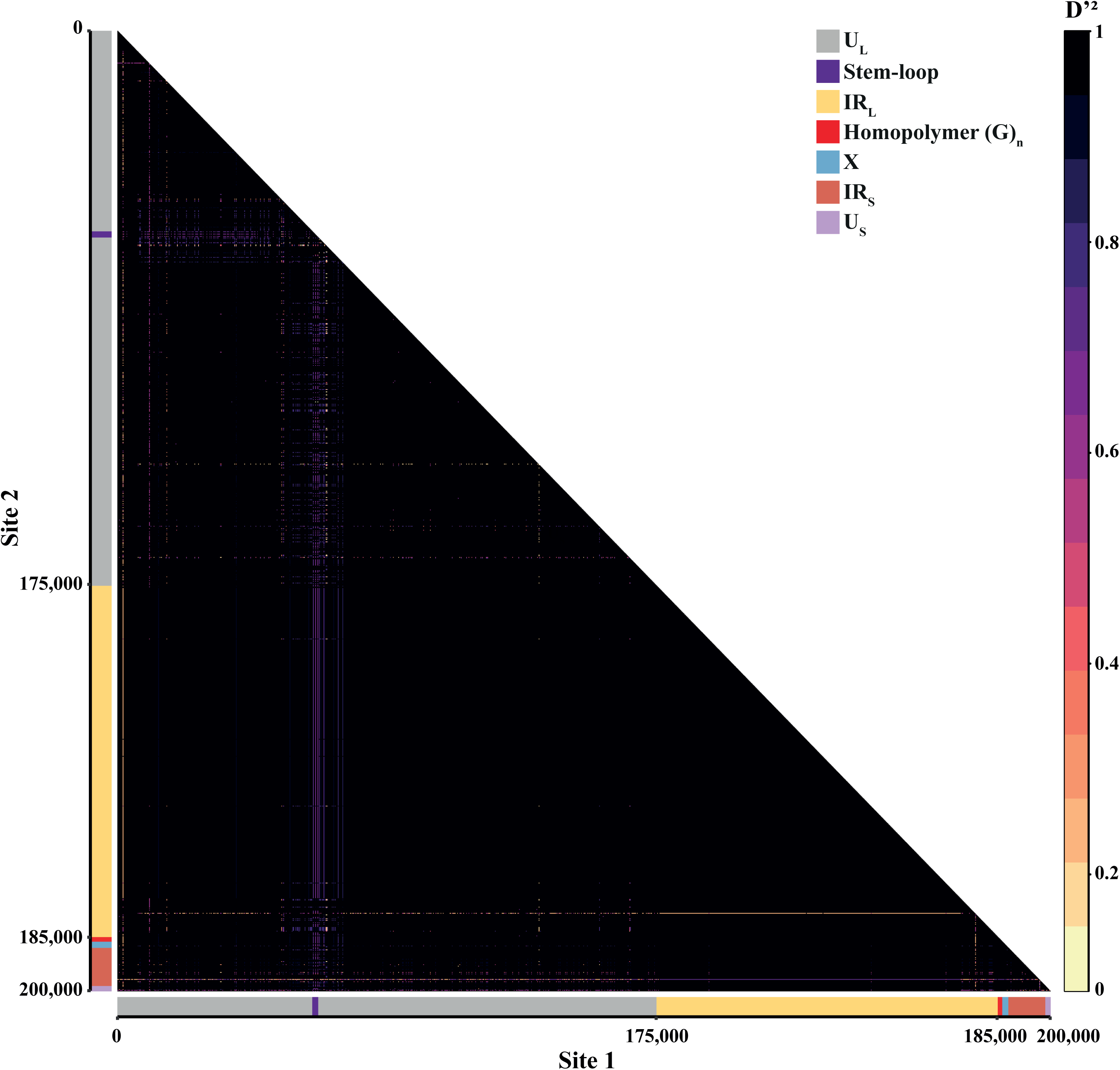
Detection of rare recombination hotspots. Linkage disequilibrium (LD) heatmap between pairs of SNPs (site 1 and 2) along the alignment of 40 *de novo* assembled OsHV-1 NR-genomes and the three OsHV-1 reference NR-genomes: OsHV-1 Ref. (NC_005881.2; Davison et al., 2005), µVarA (KY242785; Burioli et al., 2017) and µVarB (KY271630; Burioli et al., 2017). LD values vary between 0 and 1, consequently, the color scale varies respectively from beige to black. At the bottom, the architecture of OsHV-1 NR-genomes and position along the multiple alignment are represented.

### 3.6. Phylogenetic structuring of OsHV-1 reflects host-specific divergence

A maximum likelihood phylogeny was then generated using a HKY+G+I model using PhyML (Guindon et al., 2010). This tree showed that OsHV-1 genomes sampled from *M. gigas* were genetically distant from those sampled from *O. edulis* (Fig. S4). In the present study, the viral lineages were clearly structured according to host species, with two main clusters identified (Fig. S4). Within each cluster, two subclusters were identified. In the *M. gigas* cluster, subcluster 1.1 corresponded to samples related to the OsHV-1 reference genome sampled between 1994 and 2008, while subcluster 1.2 corresponded to samples related to µVar lineage sampled between 2008 and 2010. In the *O. edulis* cluster, subcluster 2.2 corresponded to samples collected between 2007 and 2009 in the Quiberon Bay, while subcluster 2.1 corresponded to the sample collected in Sweden in 2012. The clustering was found to be primarily based on host species and then on the sampling date (Fig. S4). The multiple alignment of the forty-three NR-genomes enabled us to determine the similarity percentages between every pair of genomes (Fig. S5, Table S6). To provide an overview of results, the average percentage of identities for each cluster and between clusters has been computed (Table 1). Identity percentages within clusters were close to 100%, indicating a high similarity between genomes within a cluster. In contrast, the average identity percentage between genomes varied from 91% (cluster 1.1 compared to 2.1) to 96% (cluster 2.2 compared to 2.1).

**TABLE 1.**
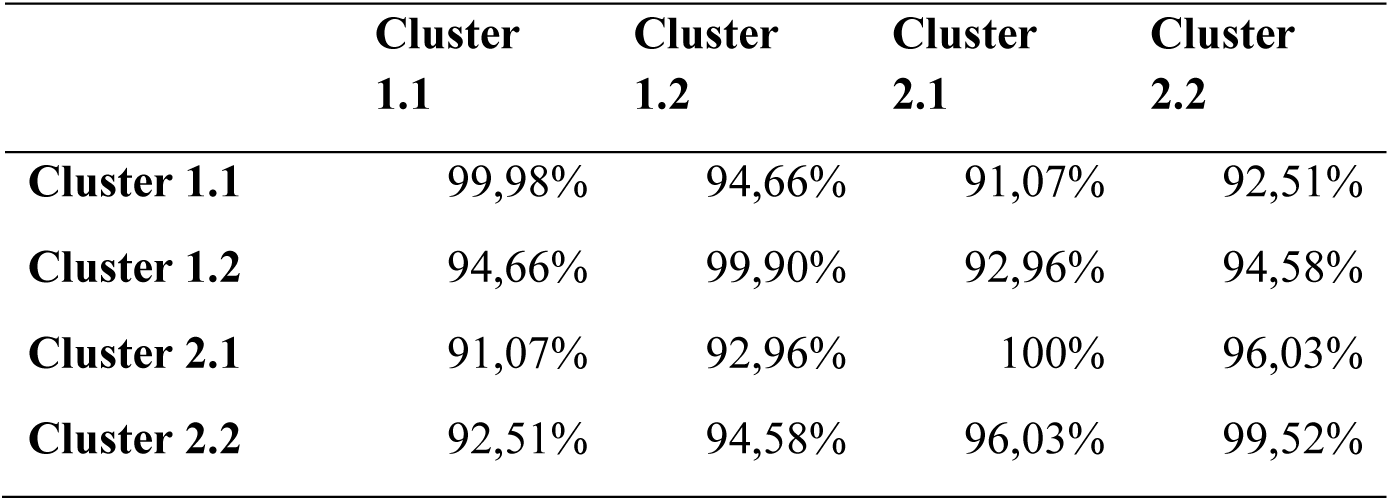
Average percentage identities within and between phylogenetic clusters based on pairwise distances computing using MAFFT.

|  | **Cluster**<br>**1.1** | **Cluster**<br>**1.2** | **Cluster**<br>**2.1** | **Cluster**<br>**2.2** |
| --- | --- | --- | --- | --- |
| **Cluster 1.1** | 99,98% | 94,66% | 91,07% | 92,51% |
| **Cluster 1.2** | 94,66% | 99,90% | 92,96% | 94,58% |
| **Cluster 2.1** | 91,07% | 92,96% | 100% | 96,03% |
| **Cluster 2.2** | 92,51% | 94,58% | 96,03% | 99,52% |

### 3.7. Consensus-based diversity analysis

Nucleotide diversity (θ_π_) and the number of segregating sites (S) varied across the different clusters analyzed. When considering all OsHV-1 genomes combined, θ_π_ was estimated at 0.003041 and S at 23161 sites, indicating a relatively high level of genetic variation. In contrast, OsHV-1 genomes assembled from *O. edulis* samples showed much lower diversity, with a θ_π_ of 0.00015 and a S of 7187 sites. OsHV-1 genomes assembled from *M. gigas* samples exhibited intermediate values, with a θ_π_ of 0.001434 and S of 11143 (Table 2).

**TABLE 2.** Nucleotide diversity θ_π_ and number of segregating sites S calculated for all OsHV-1 genomes, as well as separately for OsHV-1 genomes collected from *O. edulis* and *M. gigas* samples.

|  | All samples | *O. edulis*<br>samples | *M. gigas*<br>samples |
| --- | --- | --- | --- |
| $\Theta_{\pi}$ | 0.003041 | 0.00015 | 0.001434 |
| S | 23161 | 7187 | 11143 |

An AMOVA was performed to test the factors structuring the observed genetic differences (Table 3). The results indicate that 76.85% and 13.30% of the total genetic variance are explained by interspecies variations and the geographic origin of samples, respectively. In addition, 3.97% of the variance is attributed to differences between samples within each geographic origin, while 5.88% is explained by variations within samples.

**TABLE 3.** Analysis of Molecular Variance (AMOVA) to determine the impact of host species, origin, year of sampling and host developmental stage on the observed OsHV-1 diversity at the genomic scale.

|  | DF | SS | MS | Est. var. | % | PhiPT |
| --- | --- | --- | --- | --- | --- | --- |
| *Between Host species* | 1 | 17075 | 17075.4 | 729.8 | 76.85 % | 0.768 |
| *Between Origin within Host species* | 7 | 2934 | 419.2 | 126.3 | 13.30 % | 0.575 |
| **Between samples within Origin** | 6 | 1128 | 188.0 | 37.7 | 3.97 % | 0.403 |
| **Within samples** | 27 | 1508 | 55.9 | 55.9 | 5.88 % | 0.941 |
| **Total** | **41** | **22646** | **552.3** | **949.7** | **100.0 %** | **2.687** |

DF: degrees of freedom, SS: sum of square deviation, MS: mean square deviation, Est. var.: estimated variance, %: percentage of variance explained by the factor, PhiPT: population differentiation statistics.

The diversity analysis based on every NR consensus genome was completed with the computation of commonly used measures in population genetics, including the fixation index G_ST_ and Tajima’s D (Fig. 6). The global pairwise Nei G_ST_ was equal to 0.745, indicating the presence of different fixed alleles in the two species. To determine regions that were highly differentiated between them, G_ST_ values were computed for 1kb windows along the alignment of OsHV-1 genomes. G_ST_ values varied from 0.005 to 1 along genomes and were, on average, equal to 0.545. A threshold of G_ST_ = 0.6 was used to distinguish between highly differentiated and less-differentiated regions. Results show 34 out of 200 windows of 1kb having a G_ST_ higher than 0.6 indicating high level of genetic differentiation of these regions (Fig. 6A). Among these 34 windows, 29 were located within the U_L_ region and 5 were located in the IR_L_ region (Table S7). X, IR_S_ and U_S_ regions seemed to be less differentiated according to host species than the rest of the genome. Additionally, 100 bp windows were computed within these 1kb windows to define more precisely segregating sites that were differentiated. 87 windows of 100 bp out of 340 were detected as highly differentiated (G_ST_ > 0.6). These windows were located within 2 intergenic regions and 34 ORFs coding for proteins with mainly unknown functions (15 ORFs, see Table S7), secreted proteins (ORFs 68, 88, 5.1), proteins containing domains such as Zing-finger, Ring type proteins (ORFs 53, 97, 106), proteins with transmembrane domain (ORFs 22, 32, 50.2 54, 59, 65.1, 68, 77, 84, 88), inhibitor of apoptosis (ORFs 87, 106), ribonucleotide reductase (ORF 20), DNA replication origin binding (ORF 115), large subunit DNA-packaging terminase (ORF 109).

**Figure 6.**
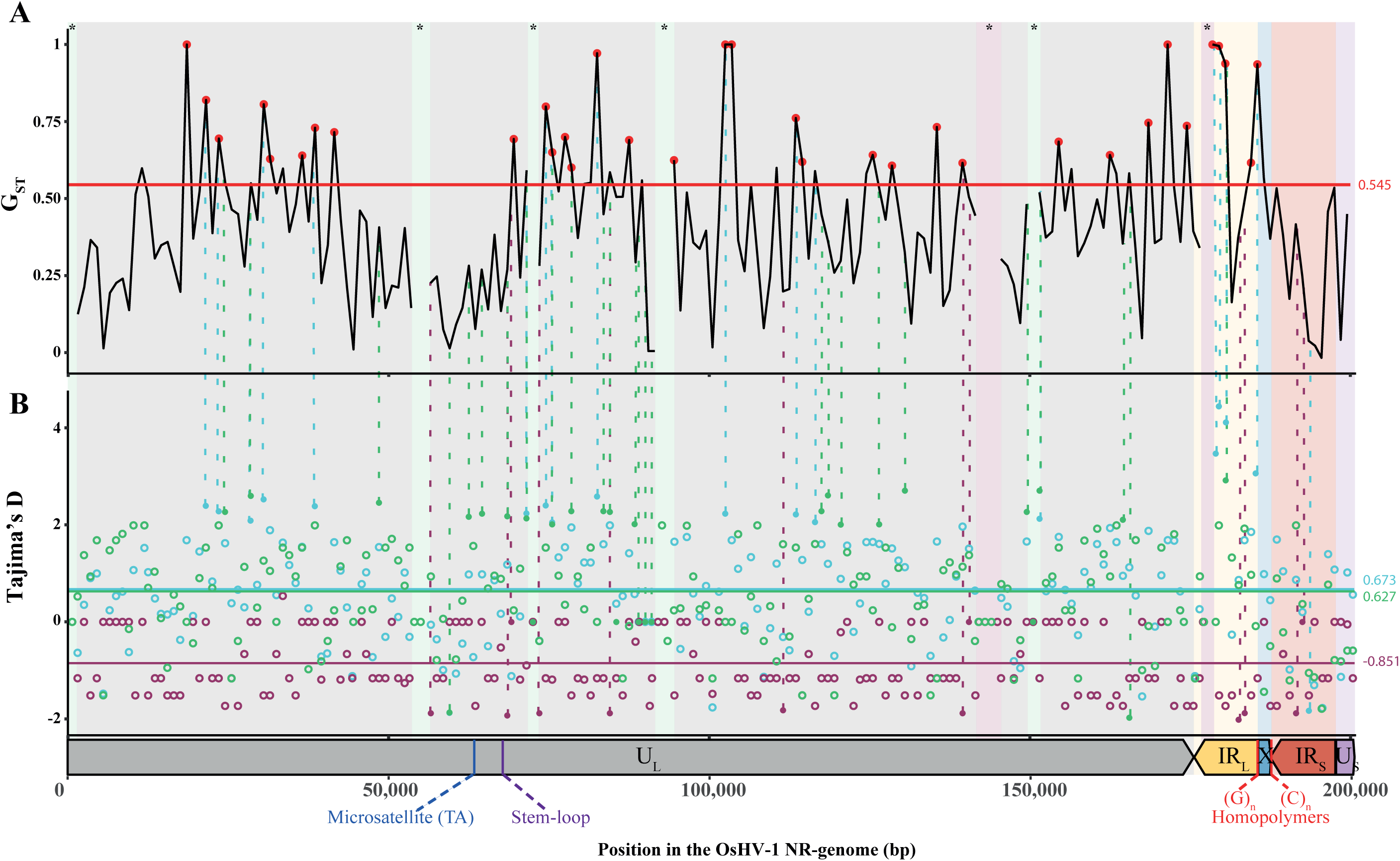
Consensus-based population genomics of OsHV-1. (A) Genetic differentiation between OsHV-1 genomes isolated from *M. gigas* and *O. edulis,* estimated using mean G_ST_ values calculated for 1kb windows along the multiple alignment. The red horizontal line at y=0.545 represents the genome-wide average G_ST_, which gives an overall estimate of the genetic differentiation of OsHV-1 between species. Red dots indicate windows with particularly high differentiation (G_ST_ > 0.6). (B) Distribution of Tajima’s D values obtained from 1kb windows across all OsHV-1 genomes (blue), for genomes sampled from *M. gigas* (green) and *O. edulis* (purple). If values of Tajima’s D were significant, dots were filled, while if not significant, dots are empty. The blue, green and purple lines represent the averaged Tajima’s D value for each genome data-set. For a better association of the two population genetic measures for a unique window, significant Tajima’s D values are linked by dotted lines to GST values. Below the graph is represented the architecture and position in the OsHV-1 NR-genomes multiple alignment. Genomic regions are reported on the two graphs above. Additionally, regions present only in *M. gigas* and *O. edulis* OsHV-1 subpopulations are represented with green and purple backgrounds respectively.

### 3.8. Contrasting evolutionary dynamics of OsHV-1 in Ostrea edulis and Magallana gigas

Tajima’s D values were calculated for 1kb windows along the alignment to identify selection patterns along OsHV-1 genomes. At the overall inter species level, most of the Tajima’s D values were positive (Fig. 6B), ranging from -1.87 to 4.44 with an average of 0.673. Later, Tajima’s D was computed for each host species. Within *O. edulis*, OsHV-1 Tajima’s D values were mostly negative, ranging from -2.02 to 0.54, with an average of -0.851. Within *M. gigas*, Tajima’s D values were predominantly positive, varying from -1.97 to 2.92, with an average of 0.627, which was comparable to the values observed for the overall population.

Tajima’s D values for the *O. edulis* lineage suggest a selective sweep or a bottleneck might have recently occurred along OsHV-1 genome. This was supported by RaISD analysis, which identified signatures of selective sweeps. In viral lineages infecting *O. edulis*, 168 regions showed µ-statistic values ranging from 1 to 8, indicative of selection. In comparison, 587 regions were affected in *M. gigas* viral lineages, with µ-statistic values from 3 to 8 (Fig. 7A). In both species, many of the regions under selection were located within ORFs encoding proteins of unknown functions. Furthermore, 8.3% to 9.9% of the regions were found in intergenic regions (Table S8). Among the annotated functional domains, the most frequently impacted were transmembrane proteins (18.5% in *O. edulis* and 17.5% in *M. gigas*), Zinc-finger RING-type domains (5.9% and 5.5% respectively), and proteins involved in viral maintenance, including DNA primase, polymerase, and genome packaging functions (Fig. 7B).

**Figure 7.**
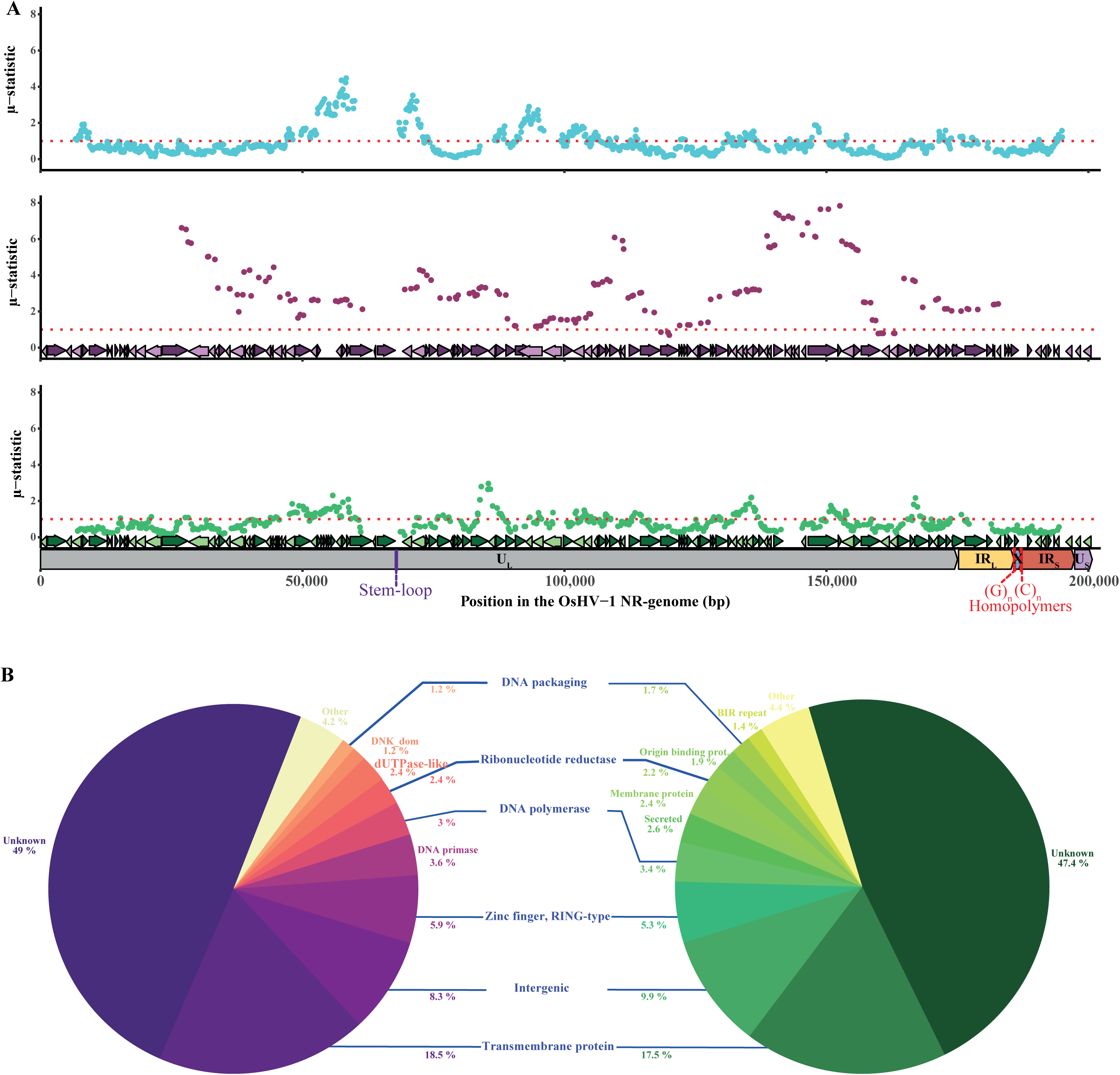
Detection of selection patterns on OsHV-1 genomes. (A) Detection of selection signatures along the multiple alignment of OsHV-1 genomes from both host species (blue points), from *O. edulis* (purple points) and from *M. gigas* (green points). Below those graphs are represented the ORFs predicted on ancestral sequence of each host species and the architecture and position in the OsHV-1 genomes multiple alignment. (B) Pie charts showing the distribution of selective sweep signals across ORFs annotated with specific functional domains for each host species. The proportions reflect the relative contribution of each functional category—such as transmembrane proteins, zinc-finger (RING-type) domains, and replication-associated genes—to the overall selection signal.

### 3.9. Ancestral states reconstruction

R² values of regression of root-to-tip genetic distance against sampling times are provided as a measure of how OsHV-1 evolution follows a clock-like pattern. According to R² values in best fitting root models, OsHV-1 harbors a sufficient temporal signal to perform molecular clock-based approaches to estimate evolutionary rates over time (Fig. S6).

The comparison of MLE values, with higher values reflecting the most appropriate models, has led to the selection of an uncorrelated relaxed molecular clock and a constant size coalescent model (Table S9).

Moreover, the evolutionary rate has been computed for the global phylogeny and for each host subcluster (Fig. S6). For the global phylogeny, the evolutionary rate has been estimated at 7.19×10^-05^ nucleotide substitutions per site per year (ns.s^-1^.y^-1^). Within the cluster from *M. gigas*, the evolutionary rate was of 5.99×10^-05^ ns.s^-1^.y^-1^ and 3.29×10^-05^ ns.s^-1^.y^-1^ for the cluster of OsHV-1 reference lineages and OsHV-1 µVar lineages respectively. In comparison, within the lineages of OsHV-1 infecting *O. edulis*, the evolutionary rate was slightly lower with a value of 1.16×10^-05^ ns.s^-1^.y^-1^.

The most recent common ancestor (MRCA) of the OsHV-1 lineages was estimated to have existed in the 1990’s (95% HPD interval [1989,1995]) and is associated with *M. gigas* (Fig. 8A). Then, the actual lineage µVar detected in France seems to have emerged around 2007 (95% HPD interval [2006,2008]). In parallel, a unique event of CST from *M. gigas* to *O. edulis* has been inferred to have occurred between 1998 (95% HPD interval [1993,2004]) and 2004 (95% HPD interval [2001,2008]) follow-up by a differentiation and a specialization of the lineages infecting *O. edulis*.

**Fig 8.**
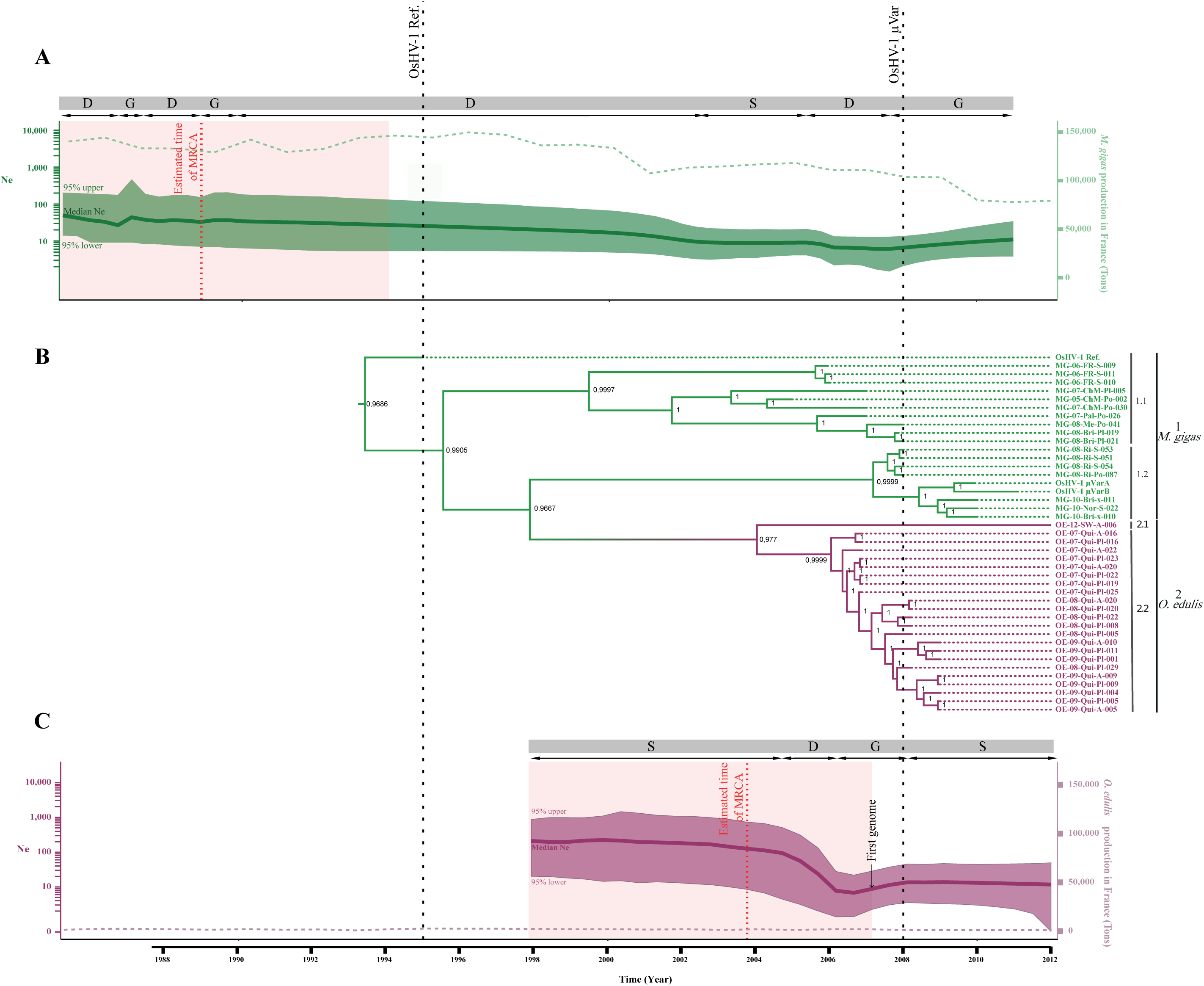
Evolution and demographic history of OsHV-1 in both oyster host species. (A) Time-scaled phylogeny of OsHV-1 genomes with ancestral state reconstruction based on a discrete host species trait. Posterior probabilities for host state assignments are indicated at each internal node. Bars and labels to the right of the tree indicate the two major viral clusters and four subclusters. (B-C) Demographic reconstructions of effective population size (Ne) over time for OsHV-1 populations infecting *M. gigas* and *O. edulis.* Solid dark lines represent inferred Ne trajectories; dashed light lines correspond to observed population sizes. Branch colors in the phylogeny and Ne trajectories denote the host species of origin (green for *M. gigas*, purple for *O. edulis*). The red dashed vertical line and red area represent, respectively, the median and the 95% highest posterior density (HPD) of the estimated time to the most recent common ancestor (MRCA). Black dashed lines represent the first detection of the OsHV-1 reference and OsHV-1 µVar lineages in France. The arrow represents the year of sampling of the first genome used in this study for both species. S: stasis, D: decline, G: growth.

### 3.10. Demographic evolution over time of OsHV-1 populations

A demographic reconstruction and estimation of the effective population size (Ne) of the viral populations over time were performed using coalescent models applied to the output trees from ancestral states reconstruction. For the viral populations of *O. edulis,* the results revealed steady decrease in Ne from 1965 to 2002 (Fig. 8C). This was followed by a sharp demographic contraction between 2002 and 2007, after which the population underwent an expansion. In recent years, however, the viral population appears to be undergoing a slow but continuous decline in size. In contrast, *M. gigas* viral lineages exhibited a fluctuating step of population size between 1982 and 1990, followed by a continuous decrease until around 2008-the year that coincides with the emergence of the µVar lineage (Fig. 8B). Following this appearance, this lineage experienced a population expansion.

## 4. Discussion

A recent genomic study revealed a potential structuring of OsHV-1 lineages according to host species raising questions about OsHV-1 host specificity, transmission cycles and cross-species transmission (CST). In this study, we compared the genomic diversity of OsHV-1 from *M. gigas* and *O. edulis* samples collected from 2005 to 2012, at both the intra- and inter-sample scales. This allowed to gain new insights into evolutionary history of OsHV-1 and the frequency of OsHV-1 CST events between the two host species.

### 4.1. OsHV-1 genome architectures

Previous studies have provided preliminary evidence that the genetic diversity of OsHV-1 is influenced by various factors, including geographical location (Delmotte-Pelletier et al., 2022; Morga-Jacquot et al., 2021), time (Burioli et al., 2016, 2017; Morga-Jacquot et al., 2021), and host species (Morga-Jacquot et al., 2021). Our analysis of structural variations identified six distinct genomic structures in our dataset, differentiated by large InDels, genomic translocations, and reversions mainly in the U_L_ and IR_L_ regions, which resulted in total size variations of approximately 6 kbp between lineages. Our findings are consistent with those previously reported on the evolution of OsHV-1 and highlighted the frequent occurrence of genomic deletions as a major driving force shaping its architecture (Morga-Jacquot et al., 2021). In addition, the total number of deletions is likely underestimated due to the use of NR-genomes to circumvent assembly challenges. The exact mechanisms underlying the generation of these structural variants are not yet understood. However, it is plausible that recombination or rearrangement events, which are common in herpesviruses, play a crucial role in their genomic architecture. Recently, long reads sequencing have demonstrated the presence of four major genomic isomers of the OsHV-1 genome that were present in stoichiometric proportions both in purified viral particle as well as in infected oyster tissues (Dotto-Maurel-Pelletier et al., 2022, Dotto-Maurel et al., 2025). In herpesviruses, and especially in the herpes simplex virus 1, the formation of isomeric genomes is closely related to the replication and recombination mechanisms that occurs early during the herpesvirus life cycle (Chaitanya, 2019; Chou & Roizman, 1985; López-Muñoz et al., 2021; Mahiet et al., 2012). During DNA replication, numerous random double-strand breaks efficiently initiate recombination events and give rise to new genomic rearrangements (Pérez-Losada et al., 2015). While these mechanisms generate isomers through segmental inversions, they can also give rise to structural variants, such as insertions, deletions, or rearrangements, which reflect deeper genomic alterations beyond isomerization alone. Importantly, these structural variants appear to be significant and not detrimental to the maintenance and replication stages of the viral life cycle. On the contrary, these variations may even be an advantage, as they potentially allow the emergence of new viral lineages and the adaptation of lineages to new hosts (Turner & Elena, 2000).

### 4.2. Population structure, diversity and host-specific differentiation of OsHV-1

Interestingly, OsHV-1 lineages sampled within the same host species were genetically very similar (with 94% and 96% genome identity for OsHV-1 sequences from *M. gigas* and *O. edulis*, respectively), whereas lineages sampled from *O. edulis* were significantly different from those found in *M. gigas* (with genome identities ranging from 91% to 94%). The genetic differentiation of OsHV-1 observed here is consistent with previous studies that have examined the genetic distances between viral genomes infecting closely related host species (Kolb & Brandt, 2020; Longdon et al., 2018). A similar phenomenon of host shift in herpes simplex virus infecting Old World monkeys, followed by co-evolution with its specific host, has been described, with a strong correlation observed between viral lineage distance and phylogenetic host distance (Kolb & Brandt, 2020). However, within the samples collected from *M. gigas*, there is also a strong clustering of lineages according to sampling date, corresponding to the OsHV-1 reference (pre-2008) and OsHV-1 µVar (post-2008) lineages. Additionally, OsHV-1 µVar lineages were not detected in the *O. edulis* samples analyzed in this study, which were collected in areas where no OsHV-1–related mortality events had been reported. These findings suggest that host species play a central role in shaping OsHV-1 evolution, with limited viral exchange across host boundaries despite ecological proximity. The absence of OsHV-1 µVar lineages in *O. edulis* may reflect host-specific barriers to infection or differences in disease ecology, which could restrict the emergence and spread of new lineages.

Among OsHV-1 clusters, contrasted differentiation and selection patterns are observed along the genome. In particular, regions coding for essential biological functions or domains involved in viral particle binding to host cells, DNA synthesis, replication and packaging, as well as trans- and membrane proteins may be critical for adapting to new host species. Indeed, viral entry proteins play a crucial role in enhancing viral fitness, and a single amino acid mutation within these proteins can alter the virus’s host range (Thakur et al., 2022; Webby et al., 2004). Furthermore, it has been shown that domains, such as envelope domains (including membrane glycoproteins and transmembrane receptors), auxiliary domains (such as Zinc-finger, RING type, dinucleoside kinase) and modulation and control domains (such as Interleukin, Interferon-regulatory factor or Zinc-finger) have been gained, duplicated, or lost during the evolution of Herpesviridae (Brito & Pinney, 2020). Many of these acquired domains allowed viruses to bind specifically to host cells and to evade or modulate the host immune system. In our study, most of the selection signals have been detected within these domains, suggesting a potential adaptation of essential proteins leading to a specialization of viral lineages to their host species.

Previous studies have not specifically investigated the intra-sample genetic diversity between infected host species but were more interested in comparing viral diversity in different organs or different individuals (*e.g.* vaccinated or unvaccinated) (Al-Khatib et al., 2022; Gu et al., 2023). However, studying within-species viral diversity is essential to better understand minor variations - i.e., variants with a frequency below 50% - as well as processes of differentiation and selection pressure. These low-frequency variants may represent emerging mutations under selection, contribute to the overall adaptability of the virus, or signal early stages of host-specific adaptation that have not yet become fixed in the population. Here, at the intra-sample scale, a clear contrast in the iSNVs detected between host species was observed. Only 92 out of 1,051 iSNVs were shared between viruses from both host species, and were located both within IR_L_ and IR_S_ and within homopolymers at the junctions IR_L_-X and X-IR_S_ of the OsHV-1 genome. The accumulation of iSNVs within IR_L_ and IR_S_ at low frequencies has already been observed (Morga-Jacquot et al., 2021). Several explanations could account for these results. First, the 7.5 kbp and 9.7 kbp regions are each duplicated within the viral genome and flank the U_L_ and U_S_ regions, respectively. Repeated regions are known, in herpesviruses, to facilitate genomic rearrangements through crossing-over events during replication (Pérez-Losada et al., 2015). These regions also contain numerous short tandem repeats that can vary in size and number between viral genomes. Some of the iSNVs detected in these areas may result from the loss or gain of such repeats.

Secondly, using a non-redundant genome to study viral nucleotide diversity may introduce bias. For instance, iSNVs located in the repeated regions (TR_L_/IR_L_ and IR_S_/TR_S_) are represented only once in the non-redundant version, potentially obscuring their full extent. Finally, sequences that are GC-rich or contain repetitive motifs, hairpin structures, or long homopolymer stretches are notoriously difficult to sequence (Kieleczawa, 2006). Repeated regions combine several of these challenging structures such as GC-rich, multiple repeats, and long homopolymer stretches, making the interpretation of iSNVs within them particularly complex. The accumulation of such variants may reflect sequencing artifacts rather than true biological variation, as polymerases used during sequencing may struggle to access these sequences due to secondary structures or DNA methylation. Sequencing platforms that avoid polymerase-based amplification, such as Nanopore sequencing, may help overcome these challenges.

The allelic association graph used to compare the co-occurrence frequency of each iSNVs among samples revealed three distinct clusters: two corresponding to *M. gigas* and one corresponding to *O. edulis* population. These results suggested that lineages present in *M. gigas* displayed a higher level of genetic diversity compared to those originating from *O. edulis.* This observation could be explained by host populations size and species ecology indeed *M. gigas* is currently the most widely cultivated oyster in the world, with enormous population sizes, both locally within oyster beds or in the various production basins. Moreover, this species is often transferred from one basin to another, particularly in France due to breeding practices (Lupo et al., 2016). In France, flat oyster populations are small and have long been concentrated in Brittany, where individuals are often moved locally between northern and southern areas during their growth. However, this does not apply to the samples used in this study, which were collected from natural, non-transferred populations. Moreover, the populations are distributed in "patches", which does not allow direct connections between the different populations and limit viral transmission (Nørgaard et al., 2019). The unexpectedly large effective population size (Ne) inferred for OsHV-1 infecting *O. edulis* could be explained by the possibility that these viral lineages also infect other, non-sampled host species, thereby increasing the overall host population size. This hypothesis is supported by evidence that the virus is known to circulate in multiple host species beyond *O. edulis* (Arzul et al., 2017; Arzul, Nicolas, et al., 2001; Arzul, Renault, & Lipart, 2001; Arzul, Renault, Lipart, et al., 2001; C.-M. Bai et al., 2019; Comps, 1988; Ren et al., 2013; Renault et al., 2001; Xia et al., 2015).

The fine-scale diversity observed within OsHV-1 reinforces findings obtained from consensus-based analyses and confirms the hypothesis that the virus is differentiated based on host species and has undergone coevolution with its respective new host following adaptation. An important avenue for future research would be to investigate the coevolution between OsHV-1 and multiple host species using phylogenetic approaches. This would involve sampling different host species from various locations to obtain a comprehensive understanding of the coevolutionary dynamics between the virus and its hosts and to give further insights into the evolutionary processes and host-virus interactions shaping the viral diversity pattern.

As previously mentioned, the selection of samples was designed to maximize heterogeneity in host-developmental stage, sampling date and origin, in order to explore the potential effects of various factors on viral diversity, while still being constrained by sample availability, particularly for flat oysters. Consequently, it is essential to consider that the variability in host developmental stage, sampling period, and geographic origin may have introduced biases into our results (Chikhi et al., 2010; Sul et al., 2018). For instance, this variability may have resulted in the overrepresentation or underrepresentation of specific viral lineages or sublineages associated with particular factors such as host developmental stage or the timing of sampling post-infection. Such biases could potentially compromise the accuracy of our representation of the viral diversity and significantly affect the reconstruction of ancestral states (Liu et al., 2022). Furthermore, the heterogeneity of our dataset introduces challenges in disentangling confounding effects among factors, making it difficult to differentiate between clusters parameters and establish clear causal relationships between variables and their impact on the viral diversity. This complexity can hinder our ability to draw definitive conclusions about the specific factors influencing viral dynamics. Moreover, comparisons between samples may be limited due to inherent differences, such as temporal dynamics in viral spread or diversity across species, host stages, sampling location or sampling years. These differences introduce additional complexities and may hinder direct comparisons between samples, potentially impacting the generalizability of our findings.

### 4.3. Rare recombination and cross-species transmission events

Cross-species transmission events have been identified as a key driver of genomic rearrangements and recombination events that facilitate host adaptation and specialization. An experimental evolution study has shown that structural rearrangements of the 3’-UTR of the Dengue virus genome were observed after 10 generations, leading to differential viral lineages within mosquitoes and human (Villordo et al., 2015). These rearrangements could potentially confer a high capacity for rapid adaptation and evolution in response to host shifts. Although our analysis of OsHV-1 identified potential recombination events, only a limited number of them were detected. Specifically, five recombination events were identified between the stem-loop regions and a microsatellite (TA)_n_ located in the U_L_ region, as well as in the homopolymer regions at the junctions of the X with repeated regions. Notably, it has been demonstrated that a bias in the detection of recombination breakpoints could occur in repetitive regions, intergenic regions, and areas with high G+C content (Lee et al., 2015). This finding provides a potential explanation for the occurrence of only five recombination events observed in our study. Moreover, the frequency of recombination events in herpesviruses is not well-known (Renner & Szpara, 2017). However, studies have demonstrated numerous recombination events between Herpes simplex virus 1 and 2 (Burrel et al., 2017; Koelle et al., 2017) as well as between strains of Human cytomegalovirus (HCMV). These recombination events were detected throughout the genome and mainly in unique long regions (Lassalle et al., 2016; Sijmons et al., 2015) as observed in our dataset. The genome of OsHV-1 may have undergone evolutionary processes that have led to these rare occurrences of recombination and potentially contributed to the adaptation of lineages to their respective hosts (Parrish et al., 2008). To fully understand the implications of the observed rearrangements and recombination events within the genome of OsHV-1, additional investigations are required. Specifically, it is crucial to examine additional OsHV-1 samples sourced from diverse geographic locations and additional host species. This would help to determine whether similar patterns of genomic rearrangement and selection exist across different populations. Additionally, conducting experimental infections studies aiming at elucidating the functional consequences of these genetic modifications would provide valuable insights into their biological significance.

OsHV-1 is a measurably evolving virus indeed at the global genomic scale, the mean evolutionary rate inferred in this study was 7.19×10^-05^ ns.s^-1^.y^-1^, a value comparable to that previously reported (Morga-Jacquot et al., 2021). Within the *M. gigas* cluster, substitution rates ranged from 5.99×10^-05^ to 3.29×10^-05^ ns.s^-1^.y^-1^, whereas within the *O. edulis* cluster, the rate was slightly lower, estimated at 1.16×10^-05^ ns.s^-1^.y^-1^. These findings are consistent with expectations, as the effective population size of *M. gigas* is substantially larger than that of *O. edulis*, which is currently in decline. Nevertheless, the evolutionary rate of OsHV-1 remains markedly higher than those estimated for members of the Herpesvirales order evolving under co-speciation, with rates of 3.0×10^-09^ ns.s^-1^.y^-1^ for avian herpesviruses (McGeoch et al., 2000), 4.4×10^-09^ ns.s^-1^.y^-1^ for herpesviruses collected only from mammals (McGeoch et al., 1995) and 4.37×10^-09^ ns.s^-1^.y^-1^ for cyprinivirus (Donohoe et al., 2021). These differences in evolutionary rates are likely driven by the intensive farming conditions of oysters and the contrasting dynamics of wild host population sizes. Oysters typically live in close proximity and at high population densities, conditions that are conducive to viral transmission. It is well established that increased transmission opportunities enhance viral replication rates, thereby accelerating viral evolution and promoting greater genetic diversity (Helmer et al., 2019; Peck & Lauring, 2018).

The analysis of ancestral traits indicate that OsHV-1 has evolved and adapted into two distinct phylogenetic cluster, one infecting *M. gigas* and the other infecting *O. edulis*. The most recent common ancestor (MRCA) of these two clusters was traced back to *M. gigas*, suggesting that this host species may have played a crucial role in the evolution and emergence of the OsHV-1 reference lineage. Although a first event of CST was inferred to have potentially led to the emergence of two highly distinct phylogenetic clusters, no subsequent CST or inter-species recombination events have been observed while the two host species *O. edulis* and *M. gigas* are living in nearby environment. As the only three other OsHV-1 genomes assembled from infected *C. farreri* ([MF509813](https://www.ncbi.nlm.nih.gov/nuccore/KP412538.1) and GQ153938.1) and *A. broughtonii* (KP412538.1) were not collected in France and are genetically distant from the OsHV-1 lineages sampled from *M. gigas* and *O. edulis* (Dotto-Maurel-Pelletier et al., 2022; Morga-Jacquot et al., 2021), they were not included in the study. Moreover, no other samples collected from different host species were available to complete the dataset. These results obtained in this study provide an initial insight into the evolutionary history of OsHV-1, but need to be supplemented by the contribution of other viral genomes sampled from different species to understand the involvement of different host species in viral diversity and infer the full range of CST events.

Taken together, our results suggest that OsHV-1 was likely introduced into Europe with *M. gigas*, from which it subsequently underwent a host shift leading to the emergence of a genetically distinct lineage infecting *O. edulis*, between 1998 and 2004. This initial CST event appears to have been followed by strong host-specific divergence, giving rise to two separate phylogenetic lineages. The evolutionary trajectory observed here reflects a scenario in which the virus, after an initial CST event, adapted to a new host and became differentiated within distinct host species, likely shaped by ecological barriers and host-specific selective pressures.

The frequency of CST occurrences is determined by the presence of facilitators or barriers encountered by the virus (Tian et al., 2022; Webby et al., 2004). CST events are particularly frequent in RNA viruses because of their rapid evolution (Parrish et al., 2008). The success of these viruses in infecting new hosts has been linked to their ability to adapt to a wide range of host species, notably by changing their virus surface proteins (Long et al., 2019) and by using different strategies to evade the host’s immune system, such as the production of non-structural proteins that interfere with the host immune system (Shao et al., 2017). Moreover, segmented genomes provide advantages for antigenic drift, antigenic shift, and recombination between strains infecting the same host, thus producing new combinations of genes that potentially favor the emergence of new viruses and increase their ability to infect and spread in new and unexpected host species (Both et al., 1983; Kim et al., 2018; Shao et al., 2017; Webster et al., 1982). By contrast, CST events in herpesviruses are extremely rare due to intrinsic genomic properties that lead to a persistent or latency stage rather than CST. Firstly, herpesviruses use specific receptors, such as Class I, II and III membrane proteins or entry glycoproteins, that are specific to the herpesvirus family to enter host cells. The envelope contains over a dozen viral glycoproteins, several of which are involved in the entry process (Krummenacher et al., 2013). In the case of OsHV-1, ten Class I membrane proteins, primarily responsible for facilitating the fusion of viral and cellular membranes, were predicted in its genome (Burioli et al., 2017; Davison et al., 2005). Notably, seven of these ten Class I proteins (ORFs 32, 54, 59, 65, 68, 77, and 88) showed differentiation based on host species and all were subject to selection, indicating a high degree of specificity of the viral genes toward particular host species. In addition, herpesviruses have co-evolved with their specific host species over at least 200 million years, leading to specific adaptations that enhance their ability to infect and persist within their natural host rather than infecting multiple hosts (Kaján et al., 2020; McGeoch et al., 2006; Waltzek et al., 2009). A herpesvirus that is adapted to one species may not be able to effectively be transmitted to or evade the immune system of a different species, reducing its ability to establish an infection (O’Connor & Sen, 2021). Furthermore, herpesviruses have a low mutation rate, because of error correcting activity of exonuclease-polymerase, which does not allow a rapid evolution and adaptation to the new host species. OsHV-1 relatively high mutation rate, may lead to a better adaptation ability to new host environment, meaning that it may be more likely to infect different oyster species or adapt to changes within the same species. It can also facilitate the evasion of host immune responses by producing variants that are less recognizable to the oyster’s immune system, allowing it to persist and cause infection more effectively.

In the specific case of OsHV-1, CST events between oysters may be potentially limited by ecological and physiological barriers. Indeed, the co-occurrence of *M. gigas* and *O. edulis* in oyster production areas remains relatively limited, being restricted to a few sites such as Brittany, Normandy and Languedoc-Roussillon, Marennes-Oléron and Arcachon. Furthermore, the phylogenic divergence and genetic distance between these two oyster species imply the existence of physiological barriers, notably due to species-specific immune responses (Danic-Tchaleu et al., 2011). In our study, selective sweep signals were detected within the ORFs coding for transmembrane and membrane proteins, as well as for other functions involved in the maintenance of the replication process and packaging of the virus suggesting functional to distinct host environments and therefore lineage-specific adaptation of OsHV-1 to distinct host species. Moreover, a previous study involving the injection of flat oysters with OsHV-1 isolated from naturally moribund Pacific oysters, resulting in lower viral loads and reduced mortality rates compared to *M. gigas* infections (López Sanmartín et al., 2016). These findings suggest the presence of specific host-virus interactions during the immune response (López Sanmartín et al., 2016). However, regarding OsHV-1 differentiation and its contrasting evolution in both species, the co-cultivation of the two oyster species *M. gigas* and *O. edulis* does not stand for a major infection hazard, but it could increase genetic recombination and CST events opportunities thus potentially promoting the emergence of variants that could be problematic.

## 5. Conclusion

In this study, our objective was to explore the diversity of OsHV-1 within oysters, between oysters, and among two oyster species. This study aimed to shed light on the evolutionary history of OsHV-1 by examining potential CST events and subsequent genetic differentiation of viral lineages. Our findings align with previous research and emphasize the significant influence of host species on the genetic diversity and differentiation of OsHV-1 clusters. Within *M. gigas* and *O. edulis*, OsHV-1 has evolved into two distinct phylogenetic clusters in France. One, known as the OsHV-1 µVar, primarily infects *M. gigas*, while the other specifically infects *O. edulis*. Our analysis revealed that viral CST events between these two species are rare.

Furthermore, our analysis identified selection signals within regions responsible for crucial biological functions and domains involved in viral particle binding, DNA synthesis, replication, and packaging. These findings suggest the possibility of OsHV-1 undergoing adaptation and specialization in response to its respective host species.

To deepen our understanding of the evolutionary processes and host-virus interactions, research should focus on investigating the coevolution between OsHV-1 and a wider range of host species using phylogenetic approaches. Considering the obtained results and the differentiation among various viral lineages, it would be prudent to improve the classification of *Malacoherpesviridae* by incorporating the observed lineages found in different host species. By conducting such studies, we can gain valuable insights into the diversification, adaptation, and coevolutionary dynamics of OsHV-1 with its various hosts.

## Data availability

Raw data as well as reconstructed genomes have been deposited on the NCBI database under Bioproject PRJNA999485 accession numbers SAMN36736021 to SAMN36736060 for future reference and accessibility (Table S1). All scripts used in this study are available in the GitLab repository: https://gitlab.ifremer.fr/asim/edulis_vs_gigas.

## CRediT authorship contribution statement

**Camille Pelletier**: Data curation, Formal analysis, Investigation, Methodology, Software, Visualization, Writing - original draft. **Germain Chevignon**: Supervision, Writing - original draft. **Nicole Faury**: Investigation. **Isabelle Arzul**: Resources. **Céline Garcia**: Resources, Funding acquisition. **Bruno Chollet**: Investigation. **Tristan Renault**: Resources. **Benjamin Morga**: Conceptualization, Supervision, Writing - original draft. **Maude Jacquot**: Conceptualization, Supervision, Writing original draft.

## Acknowledgments

The present study was supported by the French Institute for Exploitation of the Sea (Ifremer), by DGAL (French General Directorate for Food) through the National Reference Laboratory for Mollusc Diseases and by the European-Union Reference Laboratory for Mollusc Diseases, Ifremer, La Tremblade. CP was supported by grant from the Ifremer Scientific Board and the Nouvelle-Aquitaine region. We also thank the SeBiMER team from Ifremer for their technical assistance in bioinformatic. The authors acknowledge the Pôle de Calcul et de Données Marines (PCDM; https://wwz.ifremer.fr/en/Research-Technology/Research-Infrastructures/Digital-infrastructures/Computation-Centre) for providing DATARMOR computing and storage resources.

