## Supplemental Figures for "Phylogenomic evidence for host specialization and genetic divergence in OsHV-1 infecting *Magallana gigas* and *Ostrea edulis*"

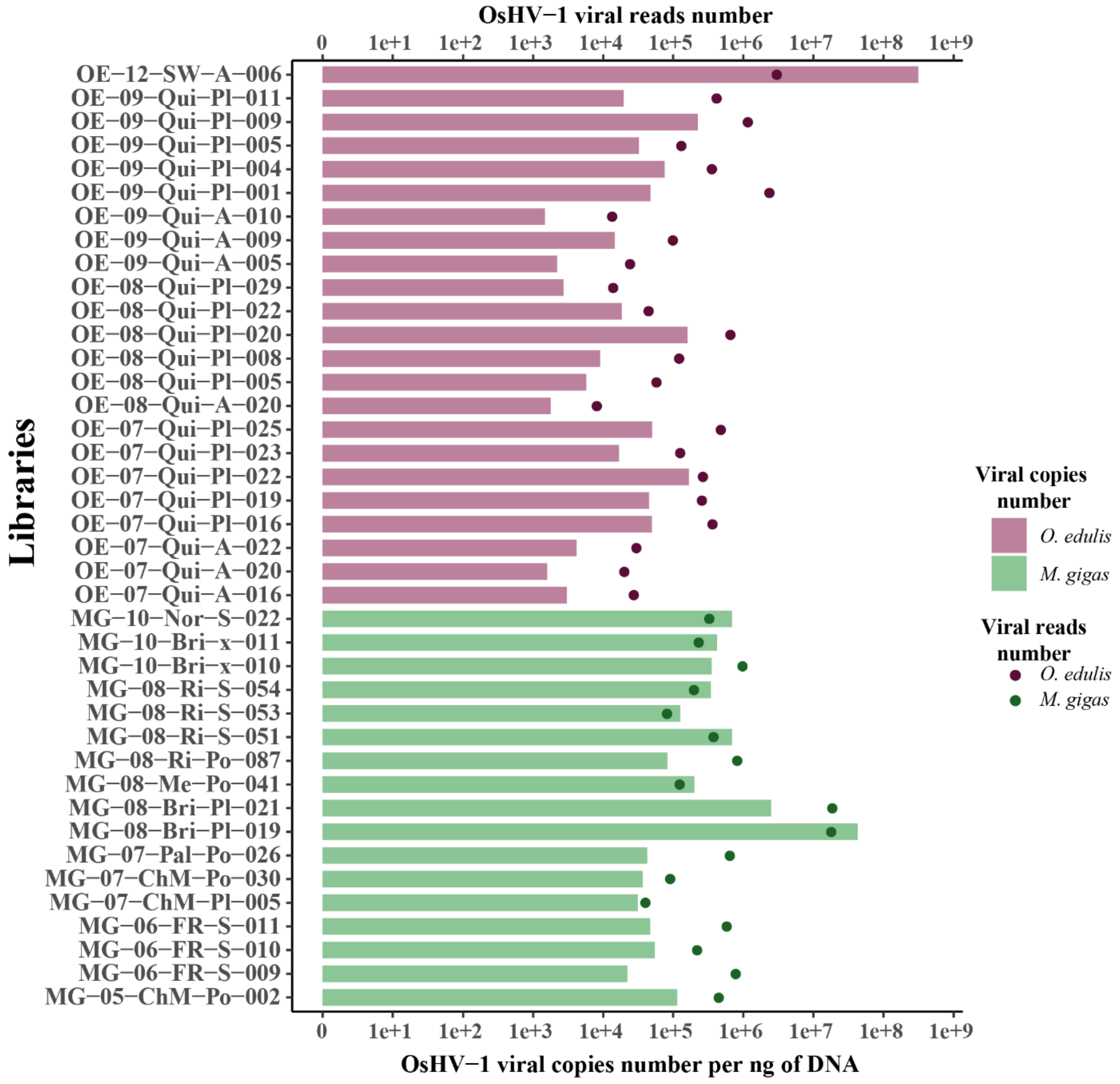


**Fig. S1: Viral copies and reads number in the libraries used in the analysis.** Bars represent the viral copies number per ng of DNA quantified by qPCR and points represent the viral reads number in each library. The color code is consistent with other figures: purple for samples from *O. edulis* and green for samples from *M. gigas*.

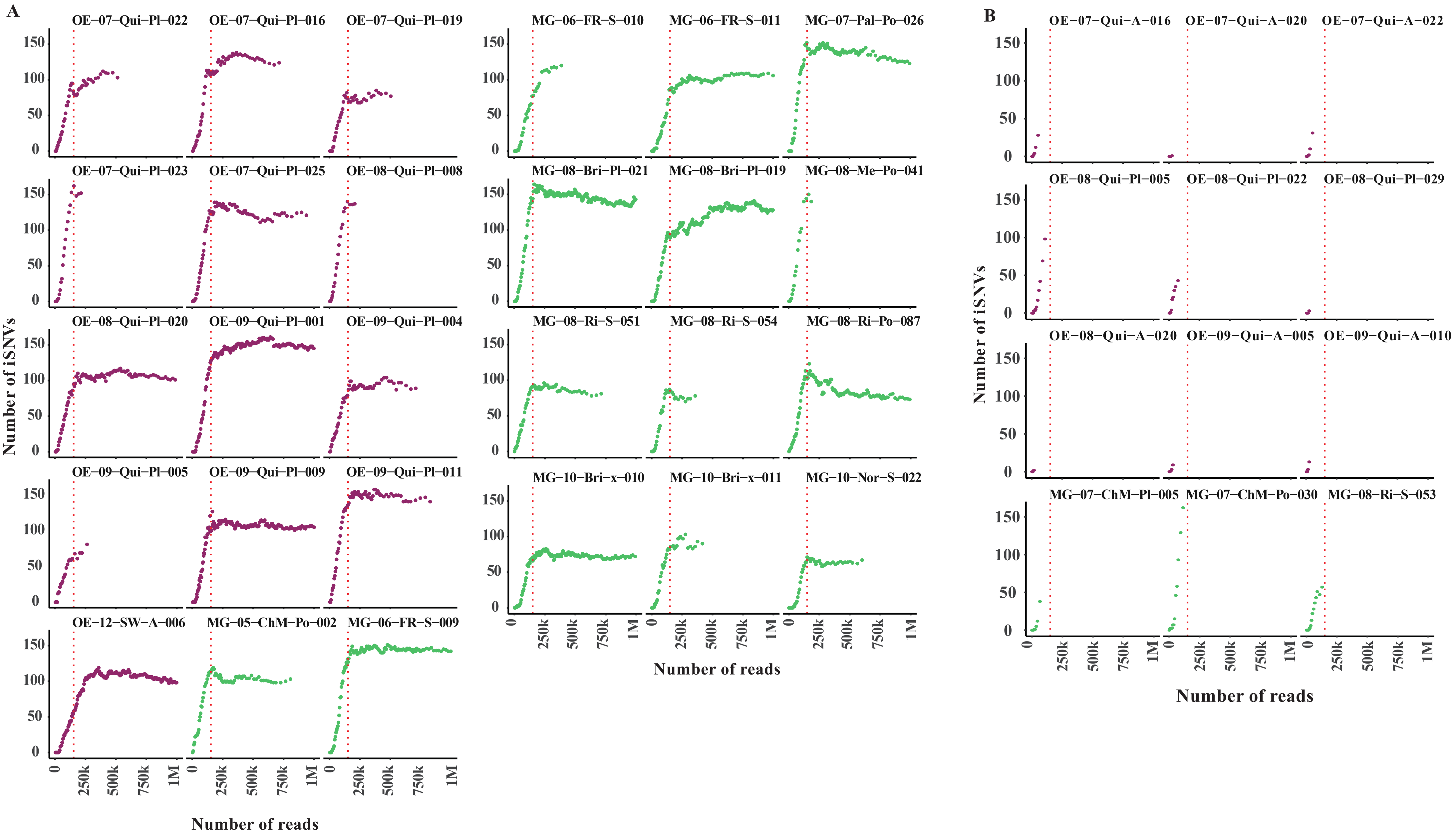
A
OE−07−Qui−Pl−022
OE−07−Qui−Pl−016
OE−07−Qui−Pl−019
OE−07−Qui−Pl−023
OE−07−Qui−Pl−025
OE−08−Qui−Pl−008
OE−08−Qui−Pl−020
OE−09−Qui−Pl−001
OE−09−Qui−Pl−004
OE−09−Qui−Pl−005
OE−09−Qui−Pl−009
OE−09−Qui−Pl−011
OE−12−SW−A−006
MG−05−ChM−Po−002
MG−06−FR−S−009
MG−06−FR−S−010
MG−06−FR−S−011
MG−07−Pal−Po−026
MG−08−Bri−Pl−021
MG−08−Bri−Pl−019
MG−08−Me−Po−041
MG−08−Ri−S−051
MG−08−Ri−S−054
MG−08−Ri−Po−087
MG−10−Bri−x−010
MG−10−Bri−x−011
MG−10−Nor−S−022
Number of iSNVs
0
50
100
150
0
250k
500k
750k
1M
Number of reads
B
OE−07−Qui−A−016
OE−07−Qui−A−020
OE−07−Qui−A−022
OE−08−Qui−Pl−005
OE−08−Qui−Pl−022
OE−08−Qui−Pl−029
OE−08−Qui−A−020
OE−09−Qui−A−005
OE−09−Qui−A−010
MG−07−ChM−Pl−005
MG−07−ChM−Po−030
MG−08−Ri−S−053
Number of iSNVs
Number of reads

**Fig. S2: Rarefaction curves showing the number of iSNVs detected by Freebayes according to the read depth in (A) libraries that contain more reads than the expected number needed to accurately detect iSNVs and (B) libraries that does not contain enough reads than the expected number needed to accurately detect iSNVs.** The red dotted lines correspond to the value at which number of iSNVs detected reach a plateau corresponding to the minimum number of reads required for effective detection of iSNVs. This threshold has been set at 150k reads.

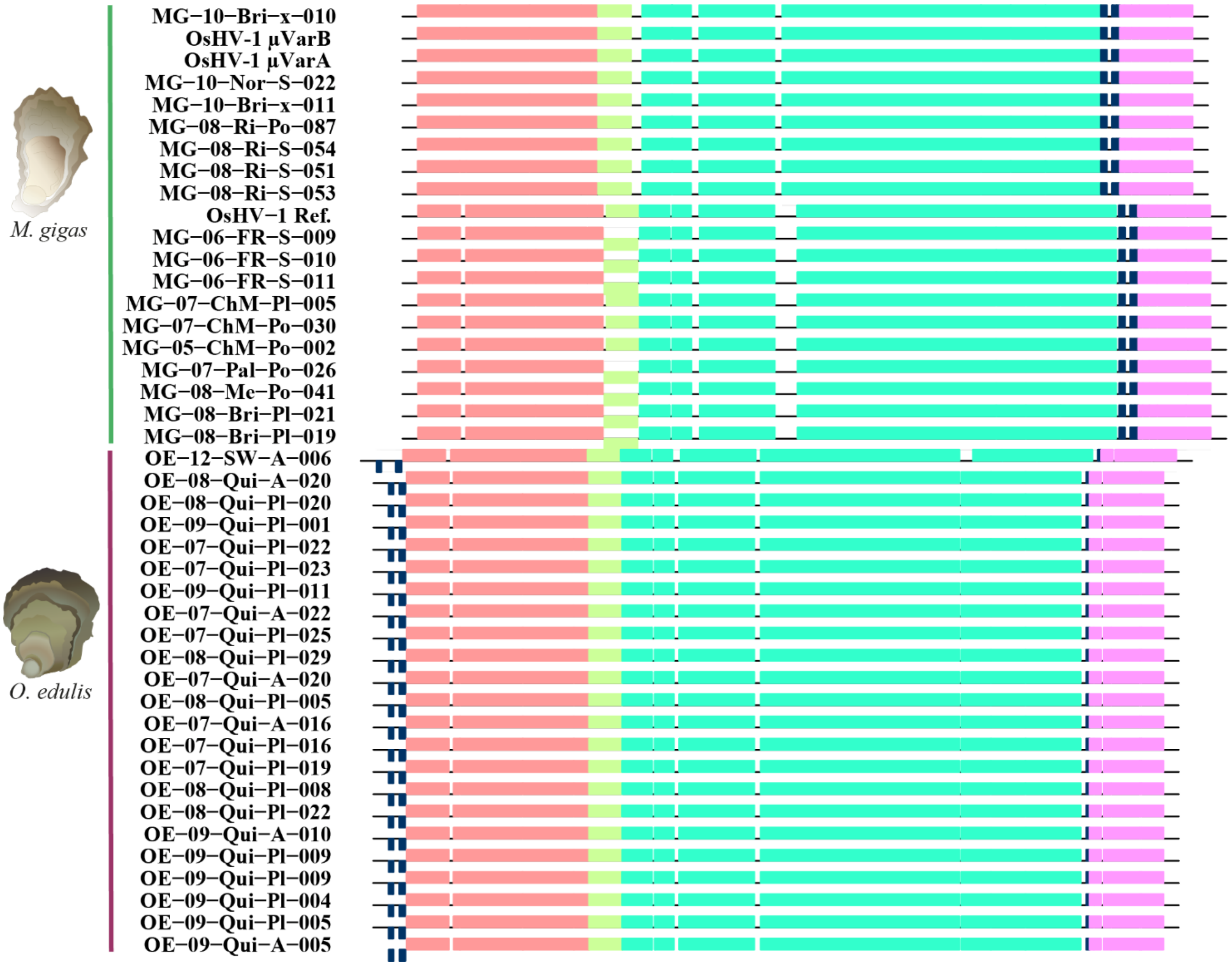


**Fig. S3: Graphical representation of the global OsHV-1 genomes multiple alignment.** The alignment contains one genome per line. Each colored block represents orthologous genomic region. White space represents gaps in the alignment. The aligned region assumes a forward orientation in relation to the initial genome sequence when a block is positioned above the central line. On the contrary, the orientation of the block is reversed when it is positioned below the center line.

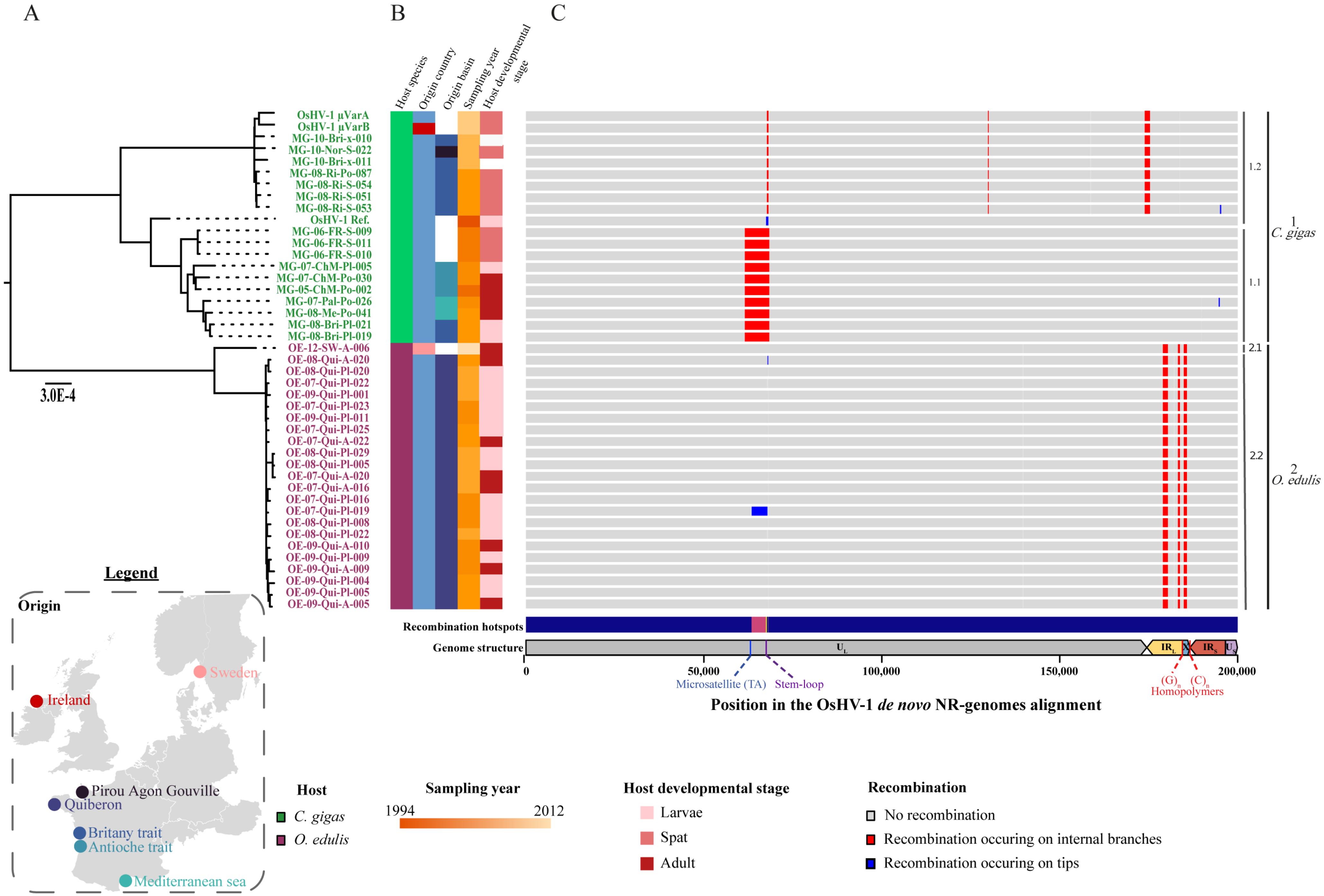

A
B
C
Host species
Origin country
Origin basin
Sampling year
Host developmental stage
OsHV-1 µVarA
OsHV-1 µVarB
MG-10-Bri-x-010
MG-10-Nor-S-022
MG-10-Bri-x-011
MG-08-Ri-Po-087
MG-08-Ri-S-054
MG-08-Ri-S-051
MG-08-Ri-S-053
OsHV-1 Ref.
MG-06-FR-S-009
MG-06-FR-S-011
MG-06-FR-S-010
MG-07-ChM-Pl-005
MG-07-ChM-Po-030
MG-05-ChM-Po-002
MG-07-Pal-Po-026
MG-08-Me-Po-041
MG-08-Bri-Pl-021
MG-08-Bri-Pl-019
OE-12-SW-A-006
OE-08-Qui-A-020
OE-08-Qui-Pl-020
OE-07-Qui-Pl-022
OE-09-Qui-Pl-001
OE-07-Qui-Pl-023
OE-09-Qui-Pl-011
OE-07-Qui-Pl-025
OE-07-Qui-A-022
OE-08-Qui-Pl-029
OE-08-Qui-Pl-005
OE-07-Qui-A-020
OE-07-Qui-A-016
OE-07-Qui-Pl-016
OE-07-Qui-Pl-019
OE-08-Qui-Pl-008
OE-08-Qui-Pl-022
OE-09-Qui-A-010
OE-09-Qui-Pl-009
OE-09-Qui-A-009
OE-09-Qui-Pl-004
OE-09-Qui-Pl-005
OE-09-Qui-A-005
3.0E-4
1.2
1.1
2.1
2.2
1
C. gigas
2
O. edulis
Recombination hotspots
Genome structure
U_L
IR_L
X
IR_S
U_S
0
50,000
100,000
150,000
200,000
Microsatellite (TA)
Stem-loop
(G)_n (C)_n
Homopolymers
Position in the OsHV-1 de novo NR-genomes alignment
Legend
Origin
Sweden
Ireland
Pirou Agon Gouville
Quiberon
Britany trait
Antioche trait
Mediterranean sea
Host
C. gigas
O. edulis
Sampling year
1994
2012
Host developmental stage
Larvae
Spat
Adult
Recombination
No recombination
Recombination occuring on internal branches
Recombination occuring on tips

**Fig. S4: Recombination events detected along OsHV-1 genomes multiple alignment.** A) Phylogenetic relationships between the 43 individuals estimated by a maximum likelihood approach. B) Metadata corresponding to OsHV-1 samples, respectively, the host species, the origin country, the origin basin, the sampling year and the host developmental stage. C) Pattern of predicted recombinations from Gubbins output. Each column relates to a base in the reference genome; each row represents an isolate in the phylogeny. Syntenic regions are represented in grey blocks. Red blocks indicate predicted recombinations occurring on an internal branch, which are therefore shared by multiple isolates through common descent. Blue blocks represent recombinations that occur on terminal branches, which are unique to individual isolates. Below the graph are represented the recombination hotspots and the architecture and position in the OsHV-1 genome multiple alignment.

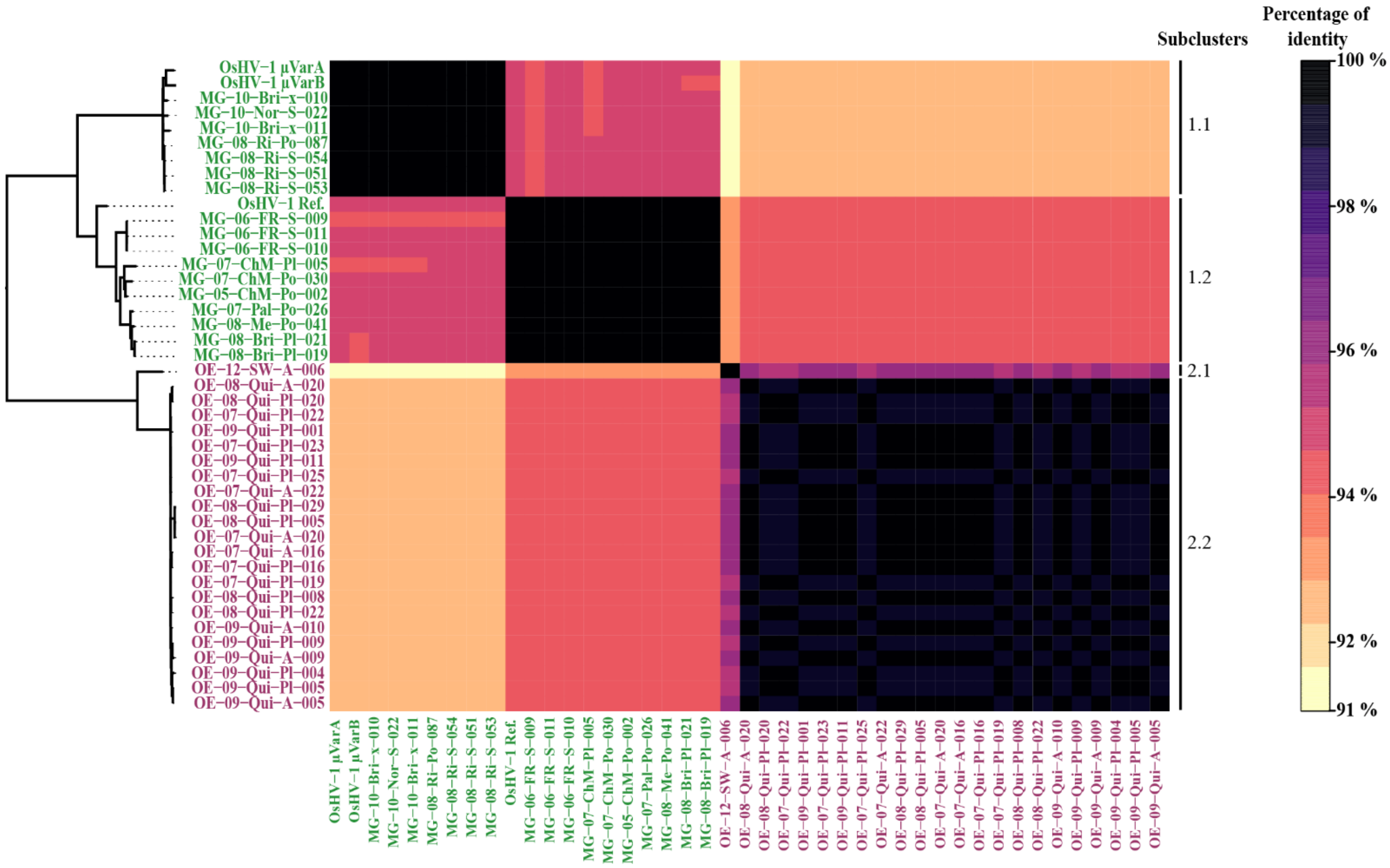


**Fig. S5: Heatmap of pairwise sequence identities comparison.** Color scale varies according to identity percentages, from black for most closely related pairs of sequences to beige for less similar sequences. On the left of the heatmap, the phylogenetic relationships between the 43 individuals are plotted. Those were estimated by a maximum likelihood approach.

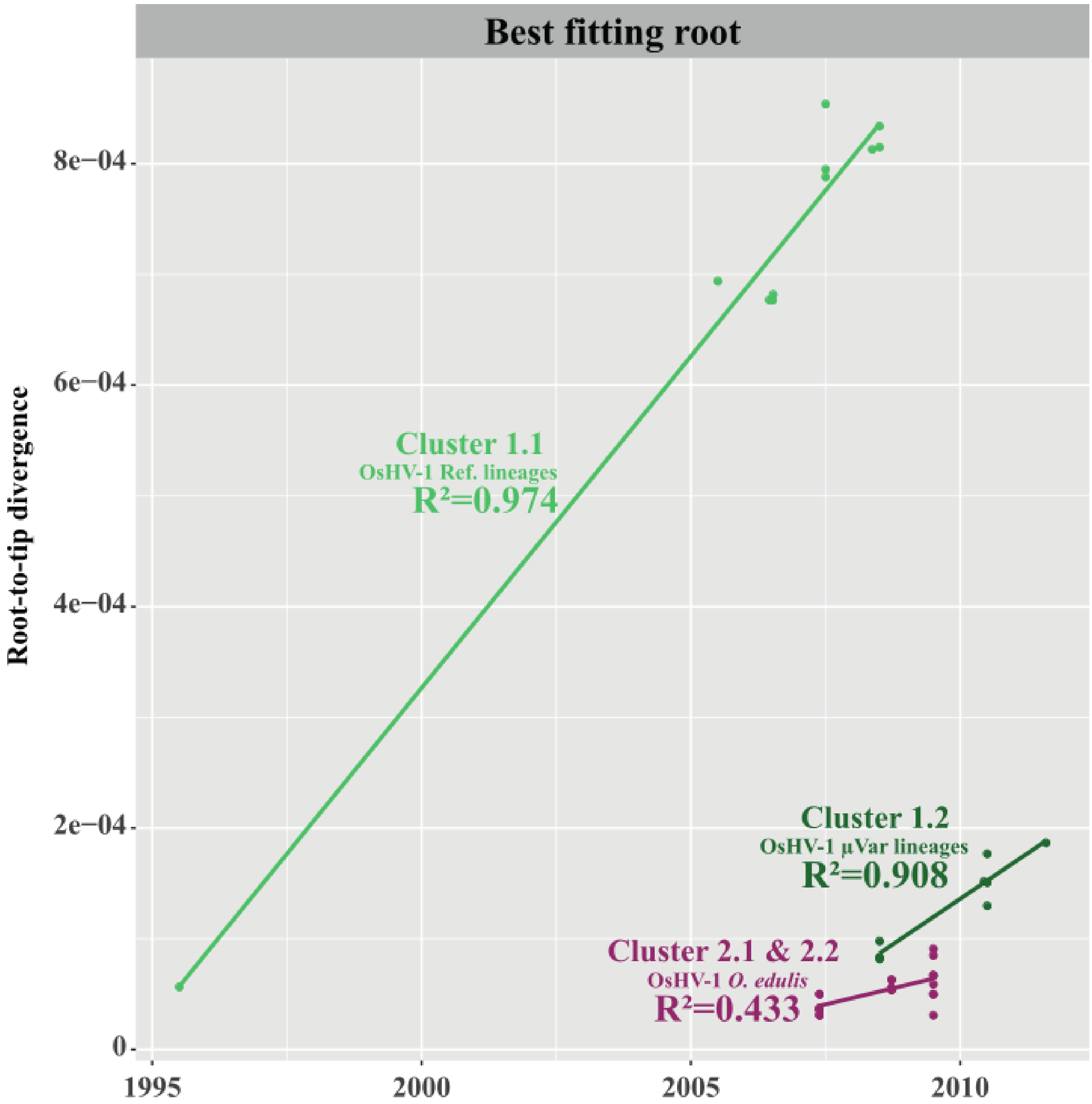


**Fig. S6: Clock-likeness of OsHV-1 genomes.** Regression of root-to-tip genetic distance against sampling time were computed for each host species based on ML tree using TempEst and the best fitting root method. Dots and lines are colored according to clusters defined on the ML tree: purple for OsHV-1 group sampled from *O. edulis,* light green and dark green for OsHV-1 group sampled from *M. gigas* reference genotypes and µVar genotypes respectively.

**Table S1. Summary of sequencing data and reads alignments statistics on OsHV-1 reference genome (NC_005881).** The 40 samples come from two host species: *Crassostrea gigas* and Ostrea edulis collected in France and Sweden between 2005 and 2012. Sequencing has been realized by different sequencing platform: [a]Ligan (CNRS UMR 8199, Lille, France); [b]Genome Quebec Company (Genome Quebec Innovation Center, McGill University, Montreal, Canada), [c]Fasteris Life Science (Plan-les-Ouates, Swiss) and [d]GenoToul (Toulouse, France).

**Table S2. Frequencies of the 985 minor iSNVs identified by variant calling analyses.**

**Table S3. Putative ORFs and associated functions impacted by iSNVs in OsHV-1 population collected from both *O. edulis* and *M. gigas*.**

**Table S4. Ratio between number of iSNVs and OsHV-1 genomic regions length.**

**Table S5. Start, end and length of recombination breakpoints detected by Gubbins and number of SNPs removed for subsequent analyses with the recombinant region.** Red values correspond to breakpoints also detected by linkage disequilibrium values.

**Table S6. Percentage of identity and number of differences between the 44 NR-genomes based on pairwise distances computing using MAFFT.** Cells are colored with a gradient between green for highly similar genomes (~99,99 % identity or less than 100 bp differences) to red for less similar genomes (~91 % identity or more than 17 kbp differences).

**Table S7. Significant genetic differentiation among OsHV-1 genomes collected in both *O. edulis* and *M. gigas*.** Initially, the mean GST values were calculated for 1-kbp windows along the OsHV-1 genomes. Subsequently, 100-bp windows were computed within each 1-kbp region that displayed substantial genetic differentiation (GST > 0.6). The table exclusively presents windows that exhibit significant GST values. For every distinct region of differentiation, information regarding the affected ORF as well as its associated function or domain are provided.

**Table S8. Summary of selective sweep events position and score detected using RaISD in both species.** Additionally, information about ORF and function or domain impacted by these events are provided.

**Table S9. Maximum likelihood estimation values used to compare the different demographic and clock models.** Values in bold correspond to the highest maximum likelihood estimation values used to select the model for ancestral state reconstruction with Beast v1.10.4 (Drummond & Rambaut, 2007). PS: Path sampling, SSS: Stepping stone sampling.
